# Centrosome amplification fine-tunes tubulin acetylation to differentially control intracellular organization

**DOI:** 10.1101/2022.10.17.512471

**Authors:** Pedro Monteiro, Bongwhan Yeon, Samuel S. Wallis, Susana A. Godinho

**Author notes:** Correspondence should be addressed to S.A.G.

## Abstract

Intracellular organelle organisation is conserved in eukaryotic cells and is primarily achieved through active transport by motor proteins along the microtubule cytoskeleton. Microtubule posttranslational modifications (PTMs) contribute to microtubule diversity and differentially regulate motor-mediated transport. Here we show that centrosome amplification induces a global change in organelle positioning towards the cell periphery and facilitates nuclear migration through confined spaces. This reorganisation requires kinesin-1 and is analogous to loss of dynein. Cells with amplified centrosomes display increased levels of acetylated tubulin, a PTM known to enhance kinesin-1 mediated transport. Depletion of α-tubulin acetyltransferase 1 (αTAT1) to block tubulin acetylation, which has no impact on control cells, rescues the displacement of centrosomes, mitochondria and vimentin, but not Golgi or endosomes. Analyses of the distribution of acetylated microtubules indicates that the polarisation of modified microtubules, rather than levels alone, plays an important role in organelle positioning. We propose that tubulin acetylation differentially impacts kinesin-1-mediated organelle displacement, suggesting that each organelle must have its own sensing and response mechanisms to ensure fine-tuning of its distribution in cells.

## Main

Eukaryotic cells display a conserved interconnected arrangement of the main cellular compartments and organelles. For metazoans, extensive changes in cell shape that occur during differentiation or migration are accompanied by organelle repositioning to maintain the functional relationship between organelles (Bornens, 2008). Thus, the ability of cells to continually adapt and respond to physiological cues requires individual organelles to be relocated. Intracellular organelle organisation is primarily achieved through active transport by motor proteins along cytoskeleton filaments (Barlan & Gelfand, 2017). The microtubule cytoskeleton, composed of αβ tubulin dimers, is polarised, with the minus-end of microtubules generally located at the centre of the cell, and plus-end located towards the cell periphery. This intrinsic polarity and distinct distribution covering most of the cytoplasm makes the microtubule cytoskeleton ideally suited to orchestrate intracellular bidirectional transport of organelles. This is important for organelle distribution and is mediated by two classes of microtubule motor proteins; minus-end directed dynein and generally plus-end directed kinesins (Barlan & Gelfand, 2017; Bryantseva & Zhapparova, 2012). Dynein and kinesin motors generate opposing pulling and pushing forces on organelles to maintain their characteristic cellular distribution, often referred to as *tug-of-war* (Sweeney & Holzbaur, 2018). Changes in the direction of transport occur when one motor wins over the other, usually in response to cellular and environmental signals (Barlan & Gelfand, 2017; Bryantseva & Zhapparova, 2012; Monzon *et al*, 2020).

Different tubulin isoforms, association with various microtubule associated proteins and tubulin posttranslational modifications (PTMs) contribute to the microtubule diversity and create different preferences for molecular motors (Janke & Magiera, 2020). Microtubules undergo numerous PTMs, including detyrosination, acetylation, phosphorylation, palmitoylation, polyglutamylation and polyglycylation (Janke & Magiera, 2020). Different tubulin isoforms and associated carboxy-terminal tail PTMs, have been shown to differentially regulate several molecular motors *in vitro* (Sirajuddin *et al*, 2014). Moreover, in cells detyrosination and acetylation can affect binding and motility of kinesin-1 motors (Balabanian *et al*, 2017; Liao & Gundersen, 1998; Reed *et al*, 2006; Tas *et al*, 2017). Thus, microtubule PTMs could play a role in organelle distribution and overall intracellular organization. Consistent with this idea, endoplasmic reticulum (ER) distribution is mediated by both tubulin acetylation and glutamylation, which regulates ER-mitochondria interactions and cytoplasm distribution, respectively (Friedman *et al*, 2010; Zheng *et al*, 2022). However, it remains unclear how these PTMs regulate the net distribution of multiple organelles and, in particular, how different organelles respond to the same modifications.

The centrosome, which is the main microtubule organizing centre in somatic cells, occupies a very characteristic position at the cell centre and in close contact with the nucleus (Bornens, 1977, 2008). This close contact requires the interaction between centrosomal microtubules and the Linker of Nucleoskeleton and Cytoskeleton (LINC) complex, composed of nesprins and SUN proteins, at the nuclear envelope (Gundersen & Worman, 2013). In addition, centrosome position at cell’s centroid is actively maintained by the radial distribution of microtubules and dynein pulling forces and also responds to anisotropic distribution of the actin network, particularly in enucleated cells (cytoplasts) (Burakov *et al*, 2003; Jimenez *et al*, 2021; Koonce *et al*, 1999). Centrosome aberrations, such as centrosome amplification, can be found in cancer cells and play direct roles in tumorigenesis (Goundiam & Basto, 2021; Nigg & Holland, 2018). Centrosome amplification can directly promote cell invasion, in part by increased microtubule nucleation (Godinho *et al*, 2014), however how these changes impact microtubule cytoskeleton and centrosome localization remains largely unknown.

In this study we discovered that inducing centrosome amplification leads to a global change in the distribution of intracellular compartments towards the cell periphery, which requires kinesin-1, suggesting it results from an imbalance of forces that favours plus-end directed motors. Cells with amplified centrosomes display increased tubulin acetylation, a known microtubule PTM involved in favouring kinesin-1 mediated motility. Systematic analyses of several intracellular compartments revealed that changes in acetylated tubulin levels differentially impacts intracellular organisation in cells. In particular, increased tubulin acetylation displaces centrosomes, mitochondria and vimentin away from the nucleus. Consistent with vimentin displacement towards cell periphery, cells with amplified centrosomes have increased nuclear deformability and migrate more proficiently through small pores. Taken together, these findings demonstrate that tubulin acetylation differentially regulates the positioning of individual intracellular cell compartments and that intracellular reorganisation could facilitate invasion in cells with amplified centrosomes.

## Results

### Centrosome amplification leads to kinesin-1 mediated centrosome displacement

In interphase cells, centrosomes localize in close proximity to the nucleus, with an average 1-2 μm distance in most cells (Rezaul *et al*, 2016). Unexpectedly, we found that induction of centrosome amplification by transiently overexpressing Polo-like kinase 4 (PLK4) using a doxycycline (DOX)- inducible system (Arnandis *et al*, 2018) (Fig EV 1A,B), led to a displacement of clustered centrosomes away from the nucleus and towards cell periphery (∼1.7 fold) in RPE-1 cells (RPE-1.iPLK4) (Fig 1A,B). This phenotype is unlikely due to unspecific effects of DOX treatment or PLK4 overexpression since overexpression of a catalytically active PLK4 truncation mutant that does not induce centrosome amplification, PLK4^1-608^(Guderian *et al*, 2010), did not lead to centrosome displacement (Fig 1B). To test if increasing cell polarization further exacerbated this phenotype, RPE-1.iPLK4 cells were embedded in a 3D collagen-I matrix that promotes cell polarization. Indeed, the distance between extra centrosomes and the nucleus was further enhanced in cells plated in 3D (∼4.1 fold) (Fig 1C,D). These results indicate that increasing centrosome numbers is sufficient to displace centrosomes away from the nucleus.

**Figure 1.**
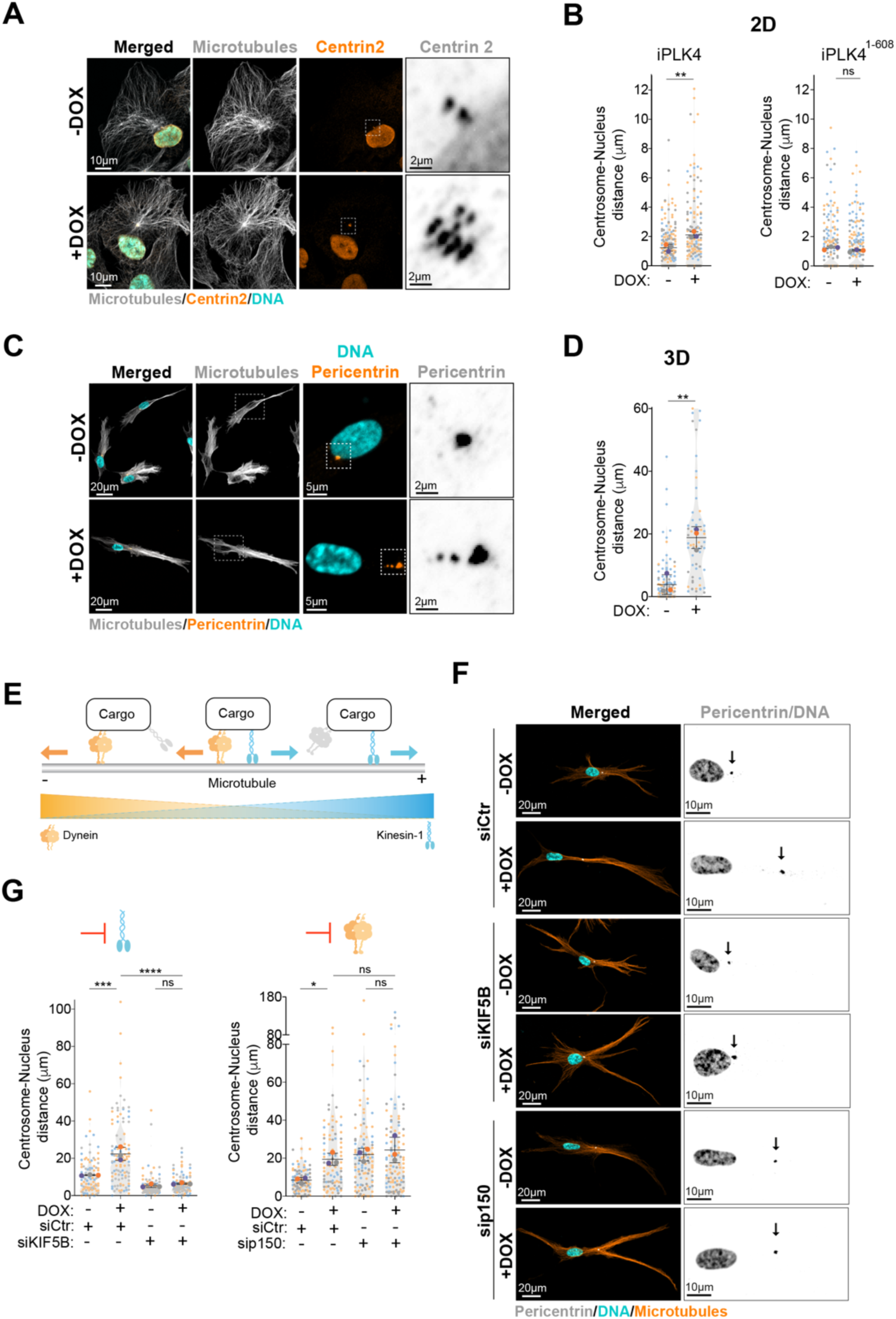
Increased centrosome displacement downstream centrosome amplification relies on microtubules and MTs-associated motor kinesin-1. **A.** Representative images of cells stained for centrosomes (Centrin2, orange), microtubules (α-tubulin, grey) and DNA (Hoechst, cyan). Scale bar: 10 µm; inset scale bar: 2 µm. **B.** Quantification of centrosome-nucleus distance in cells after induction of PLK4 (Left panel; n(-DOX)=174; n(+DOX)=162) or PLK4^1-608^ overexpression (Right panel; n(-DOX)=192; n(+DOX)=203). **C.** Representative images of cells stained for centrosomes (Pericentrin, orange), microtubules (α-tubulin, grey) and DNA (Hoechst, cyan). Scale bar: 20 µm; inset DNA/Pericentrin scale bar: 5 µm; inset Pericentrin scale bar: 2 µm. **D.** Quantification of centrosome-nucleus distance (n(-DOX)=100; n(+DOX)=62). **E.** Scheme recapitulating the balance forces mediated by kinesin-1 (blue) and dynein (orange) along microtubules. **F.** Representative images of cells stained for centrosomes (Pericentrin, grey), microtubules (α-tubulin, orange) and DNA (Hoechst, cyan) treated with siRNA control (Ctr), siRNA KIF5B or siRNA p150^glued^. Black arrows indicate the position of the centrosome(s). Scale bar: 20 µm; inset scale bar: 10 µm. **G.** Left panel, Quantification of centrosome-nucleus distance upon KIF5B depletion (number of cells: n(-DOX siCtr)=96; n(+DOX siCtr)=108; n(-DOX siKIF5B)=111; n(+DOX siKIF5B)=91); Right panel, Quantification of centrosome-nucleus distance upon p150 depletion (n(-DOX siCtr)=102; n(+DOX siCtr)=96; n(-DOX sip150)=115; n(+DOX sip150)=117). For all graphics error bars represent mean +/- SD from three independent experiments. \**p* < *0.05,* \*\**p* < *0.01, ***p* < *0.001, ****p* < *0.0001,* ns = not significant (*p* > *0.05*). The following statistic were applied: unpaired *t* test for graphs in B and D and one-way ANOVA with Tukey’s post hoc test for graphs in G.

Microtubule depolymerization by nocodazole led to a small increase in centrosome-nucleus distance, which is consistent with loss of centrosome-nucleus attachment (Salpingidou *et al*, 2007). However, no further increase was observed in cells with amplified centrosomes, suggesting that microtubules play a key role in this process (Fig EV1C,D). By contrast, latrunculin-A-mediated F-actin depolymerization did not alter centrosome-nucleus distance (Fig EV1C,D). Organelle positioning is often dictated by a balance of forces mediated by minus-end directed dynein and plus-end directed kinesin-1 motors (Belyy *et al*, 2016; Hancock, 2014) (Fig 1E). Therefore, we asked whether the displacement of centrosomes towards the cell periphery was due to imbalanced forces that favored kinesin-1. To test this, we depleted the ubiquitously expressed kinesin-1 Kinesin Family Member 5B (KIF5B) by siRNA in cells plated in 3D collagen-I matrices. We found that upon KIF5B depletion, supernumerary centrosomes remained closely associated with the nucleus, suggesting that pushing forces on the centrosomes are mediated by kinesin-1 (Figs 1F,G and EV1E). Consistent with dynein’s role in counteracting kinesin-1 pushing forces to maintain centrosome positioning (Splinter *et al*, 2010; Stiff *et al*, 2020), inhibition of dynein by depleting the p150^glued^ subunit of the dynactin complex led to similar centrosome displacement in control cells. However, depletion of p150^glued^ had no impact on centrosome displacement in cells with amplified centrosomes (Fig 1F,G and EV1E). To further evaluate if an imbalance of forces was responsible for centrosome displacement, we disrupted the interaction between centrosomal microtubules and the nuclear envelope to assess displacement in conditions where centrosomes were not constrained by its interaction with the nucleus. To do so, a dominant-negative KASH domain of nesprin-2 (^GFP^KASH2) that prevents the interaction of nesprins with the SUN proteins was overexpressed in RPE-1.iPLK4 cells (Fig EV1F,G). Overexpression of ^GFP^KASH-DL, which does not disrupt MTs association with the nuclear envelope, was used as control. We postulated that if cells with amplified centrosomes have increased kinesin-1 pushing forces, we would observe greater centrosome displacement towards the leading edge when centrosomal microtubules are not anchored to the nuclear envelope. As expected (Luxton *et al*, 2010), ^GFP^KASH2 overexpression led to centrosome displacement in control cells (Fig EV1H,I). Strikingly, once centrosomes were not constrained by their interaction with the nuclear envelope, a greater displacement towards the cell periphery was observed in cells with amplified centrosomes (∼4.5 fold) (Fig EV 1H,I). Taken together, these results demonstrate that unbalanced forces that favor kinesin-1 mediates centrosome displacement in cells with amplified centrosomes.

### Centrosome amplification leads to global intracellular reorganization

Since other organelles and cellular components rely on dynein-kinesin balance for their positioning (Barlan & Gelfand, 2017), we next investigated if centrosome amplification played a global role in organelle positioning in cells plated in 2D and 3D collagen-I matrices. Using the early endosomal antigen 1 (EEA1) marker, we assessed the distribution of early endosomes in cells with normal (-DOX) and amplified (+DOX) centrosomes. Indeed, similar to centrosomes, the distance between the nucleus and endosomes increased in cells with amplified centrosomes, but not in cells overexpressing PLK4^1-608^ (Fig 2A-C), indicating a dispersion towards the cell periphery similar to p150^glued^ depletion in control cells (Marchesin *et al*, 2015) (Figs 2D,E and EV2A). Furthermore, depletion of KIF5B in cells with amplified centrosomes resulted in endosomes repositioning near the nucleus (Figs 2D,E and EV2A). Because the intermediate filament vimentin also relies on kinesin-1 to be transported towards the leading edge (Gyoeva & Gelfand, 1991; Leduc & Etienne-Manneville, 2017; Liao & Gundersen, 1998), we assessed its distribution ratio, where a ratio>1 indicates a dispersal towards the leading edge (Leduc & Etienne-Manneville, 2017). While in control cells (-DOX) and in cells overexpressing PLK4^1-608^ vimentin remains mostly around the nucleus, in cells with amplified centrosomes (+DOX), vimentin is displaced to the cell periphery (Fig 2F-H). Dispersion of vimentin in these cells also required KIF5B, and depletion of p150^glued^ caused a similar displacement in control cells (Fig 2I,J). Additionally, we observed that both the mitochondria (labelled with MitoTracker) and the Golgi (using GM130 as marker) were also displaced in cells with amplified centrosomes in 2D and 3D cultures, but not in cells overexpressing PLK4^1-608^ (Fig EV2B-I). Interestingly, nucleus-centrosome distance seems to be most sensitive to increased cell polarization and augmented organelle displacement was not always observed in highly polarized cells plated in 3D (Fig EV2J). Taken together, these data indicate an unprecedented role for centrosome amplification in organelle organization, a process that is dependent on the kinesin-1 KIF5B.

**Figure 2.**
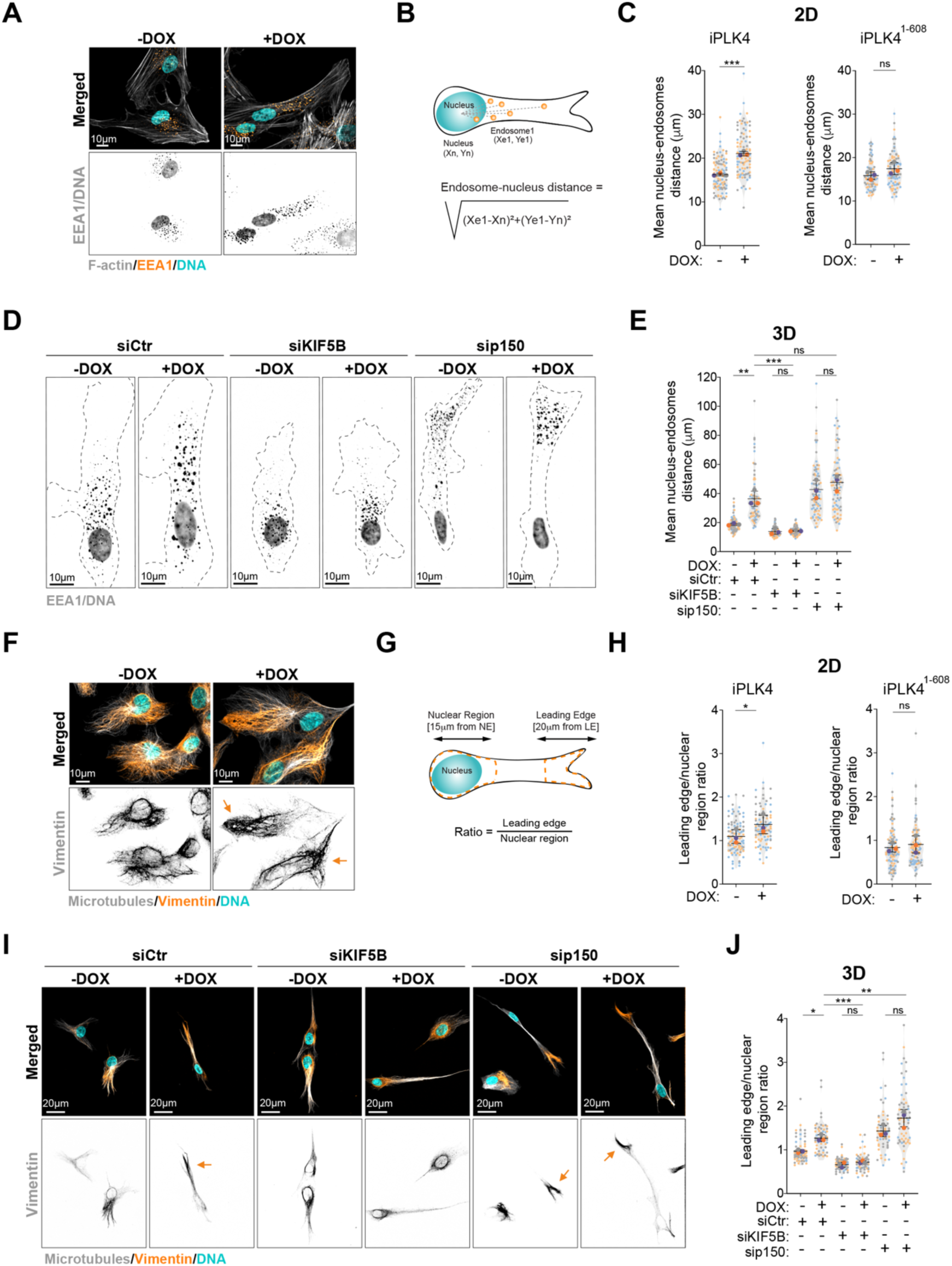
Kinesin-1 mediates endosomes and intermediate filaments displacement in cells with extra centrosomes. **A.** Representative images of cells stained for early endosomes (EEA1, orange), F-actin filaments (phalloidin, grey) and DNA (Hoechst, cyan). Scale bar: 10 µm. **B.** Representing scheme of nucleus-endosomes distance quantification. **C.** Quantification of nucleus-endosomes distance upon of PLK4 (n(-DOX)=103; n(+DOX)=100) or PLK4^1-608^ overexpression (Right panel; n(-DOX)=79; n(+DOX)=83). **D.** Representative images of cells stained for early endosomes (EEA1, grey) and DNA (Hoechst, grey). Dark dotted line represents cell contour. Scale bar: 10 µm. **E.** Quantification of nucleus-endosomes distance upon depletion of KIF5B and p150 (n(-DOX siCtr)=84; n(+DOX siCtr)=81; n(-DOX siKIF5B)=84; n(+DOX siKIF5B)=83; n(-DOX sip150)=82; n(+DOX sip150)=83). **F.** Representative images of cells stained for vimentin (orange), microtubules (α-tubulin, grey) and DNA (Hoechst, cyan). Orange arrows indicate the displacement of vimentin towards cell periphery. Scale bar: 10 µm. **G.** Representative scheme of vimentin displacement quantification. **H.** Quantification of vimentin leading edge/nuclear ratio upon PLK4 (n(-DOX)=102; n(-DOX+)=79) or PLK4^1-608^ overexpression (Right panel; n(-DOX)=104; n(+DOX)=101). **I.** Representative images of cells stained for vimentin (orange), microtubules (α-tubulin, grey) and DNA (Hoechst, cyan) upon depletion of KIF5B and p150. Orange arrows indicate the displacement of vimentin towards cell periphery. Scale bar: 20 µm. **J.** Quantification of vimentin leading edge/nuclear ratio (n(-DOX siCtr)=84; n(+DOX siCtr)=81; n(-DOX siKIF5B)=84; n(+DOX siKIF5B)=83; n(-DOX sip150)=82; n(+DOX sip150)). For all graphics error bars represent mean +/- SD from three independent experiments. \**p* < *0.05,* \*\**p* < *0.01, ***p* < *0.001,* ns = not significant (*p* > *0.05*). The following statistic were applied: unpaired *t* test for graphs in C and H and one-way ANOVA with Tukey’s post hoc test for graphs in E and J.

### Cells with extra centrosomes exhibit increased tubulin acetylation levels

Tubulin acetylation has been shown to positively influence kinesin-1 transport (Reed *et al*., 2006; Tas *et al*., 2017). Thus, we tested whether organelle displacement observed in cells with amplified centrosomes was driven by changes in tubulin acetylation. We first assessed the levels of tubulin acetylation by immunofluorescence in single cells and found that cells with extra centrosomes have a ∼2 fold increase in tubulin acetylation (Fig 3A,B). By contrast, overexpression of PLK4^1-608^ had no impact on the levels of tubulin acetylation (Fig EV3A). Cells with amplified centrosomes showed a marked increase in tubulin acetylation levels throughout the cell and near the leading edge compared to control cells (Figs 3C,D and EV3B). Tubulin acetylation has been previously associated with long-lived nocodazole-resistant microtubules and proposed to protect microtubules against mechanical ageing (Portran *et al*, 2017; Xu *et al*, 2017). We found that cells with extra centrosomes retain a significantly increased population of nocodazole-resistant that are acetylated, suggesting that increase tubulin acetylation could be a consequence of microtubule stabilization in these cells (Fig 3I,J).

**Figure 3.**
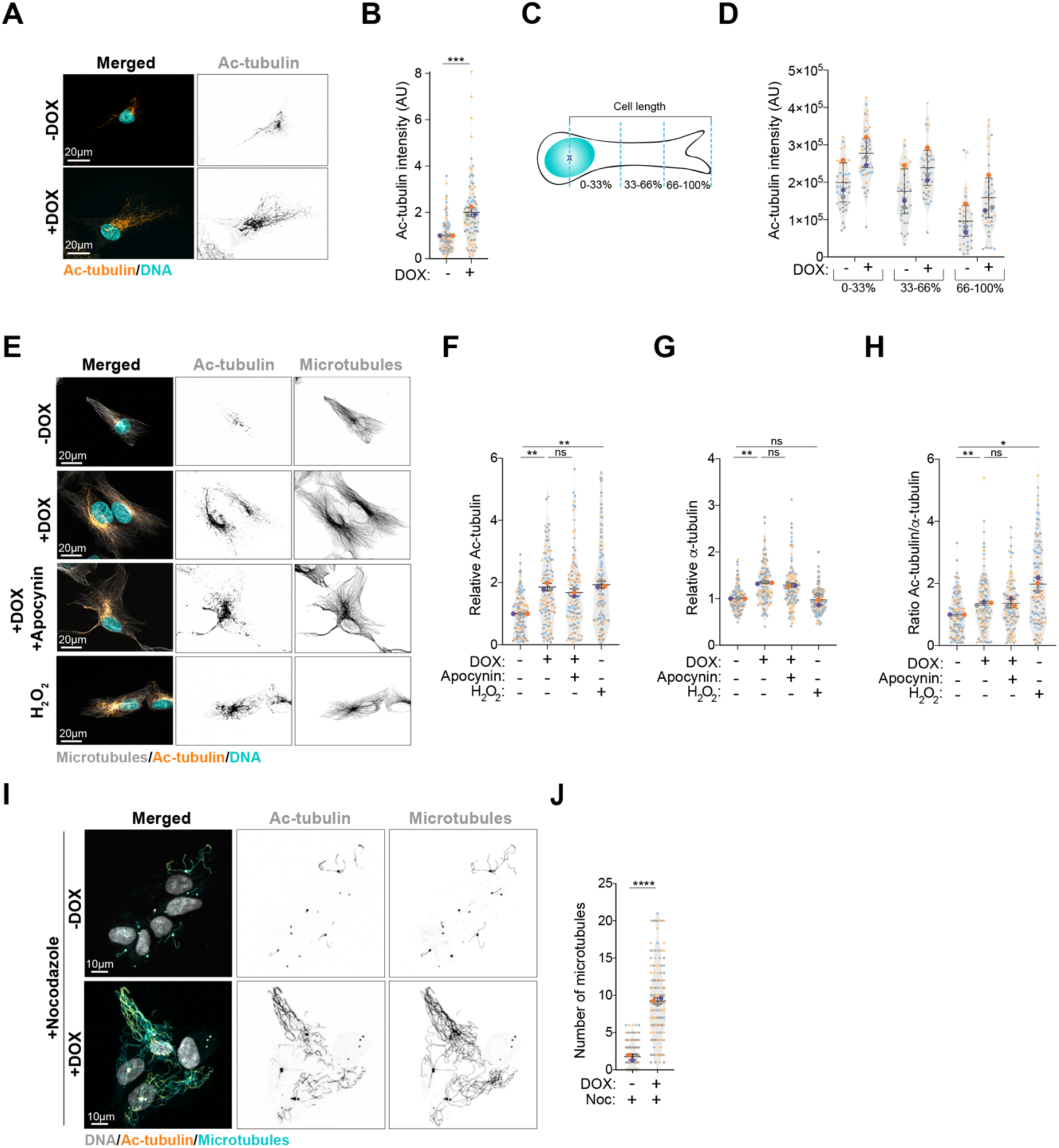
Tubulin acetylation is enhanced in cells with extra centrosomes. **A.** Representative images of cells stained for acetylated tubulin (Ac-tubulin, orange) and DNA (Hoechst, cyan). Scale bar: 20 µm. **B.** Quantification of acetylated tubulin fluorescence intensity (n(-DOX)=98; n(+DOX)=89). **C.** Representative scheme of the quantification of intracellular distribution of acetylated tubulin. **D.** Quantification of intracellular distribution of acetylated tubulin across the length of the cell (n(-DOX)=37; n(+DOX)=42). **E.** Representative images of cells stained for microtubules (α-tubulin, grey), acetylated tubulin (Ac-tubulin, orange) and DNA (Hoechst, cyan) treated with Apocynin (0.5mM) or H2O2 (75µM). Scale bar: 20 µm. **F-H,** Quantification of acetylated tubulin (**F**) and total α-tubulin (**G**) fluorescence intensity (n(-DOX)=118; n(+DOX)=110; n(+DOX Apocynin)=106; n(-DOX H2O2)=118). **H**. Ratio of acetylated tubulin to total α-tubulin. **I.** Representative images of cells stained for microtubules (α-tubulin, cyan), tubulin acetylation (Ac-tubulin, orange) and DNA (Hoechst, grey) upon nocodazole treatment (Noc, 2µM). Scale bar: 10 µm. **J.** Quantification of microtubule numbers (n(-DOX Noc)=185; n(+DOX Noc)=149). For all graphics error bars represent mean +/- SD from three independent experiments. \**p* < *0.05, ***p* < *0.001, ****p* < *0.0001,* ns = not significant (*p* > *0.05*). The following statistic were applied: unpaired *t* test for graphs in B and J. For graphs F, G, and K one sample *t* test was used for comparisons with normalised -DOX condition (using a hypothetical mean of 1) and unpaired *t* test to compare +DOX and +DOX + Apocynin conditions.

To understand the source of increased tubulin acetylation in cells with extra centrosomes, we first assessed the role of reactive oxygen species (ROS). We have previously shown that cells with amplified centrosomes have increased levels of intracellular ROS (Adams *et al*, 2021; Arnandis *et al*., 2018) and it has been recently demonstrated that hydrogen peroxide (H_2_O_2_) can damage the microtubule lattice, resulting in increased tubulin acetylation (Goldblum *et al*, 2021). We confirmed that RPE-1.iPLK4 cells with extra centrosomes (+DOX) displayed higher ROS levels, which can be blocked by treating cells with the broad NADPH oxidase inhibitor Apocynin (Fig EV3E,F). Interestingly, blocking ROS production in cells with amplified centrosomes did not prevent increased tubulin acetylation (Fig 3E,F). Because centrosome amplification can enhance microtubule nucleation (Godinho *et al*., 2014), we next tested if increased total tubulin could account for the higher levels of acetylated tubulin in these cells. Quantification of α-tubulin immunofluorescence intensity demonstrated that the presence of extra centrosomes leads to increased total tubulin in steady-state cells (Fig 3G). Normalising tubulin acetylation to total tubulin almost completely equalized the ratio of acetylated tubulin in cells with and without amplified centrosomes, although small differences can still be observed (Fig 3H). Could the increased tubulin levels be a consequence of microtubule stabilization? This is unlikely to be the case since acute H_2_O_2_ treatment increased acetylated tubulin levels without affecting total tubulin levels (Fig 3F-H). These results demonstrate that cells with extra centrosomes have increased acetylated microtubules at least in part due to higher levels of total tubulin. It is possible that in addition to damaged sites, microtubule nucleation sites, such as centrosomes, could provide an opportunity for αTAT1 to access the microtubule lumen where α-tubulin gets acetylated. This could provide an explanation for both the accumulation of acetylated tubulin around the centrosome and its increase in response to increased microtubule nucleation.

### Acetylated tubulin differentially regulates intracellular reorganization

We next assessed whether tubulin acetylation is involved in the displacement of intracellular compartments by targeting the main tubulin acetyltransferase in mammalian cells, αTAT1, which acetylates lysine 40 (K40) on α−tubulin (Akella *et al*, 2010; Shida *et al*, 2010). Using two independent siRNAs (#5 and #9) against αTAT1, we greatly reduced the levels of αTAT1 mRNA and acetylated tubulin (Fig EV4A,B). Depletion of αTAT1 rescued centrosome displacement in cells with amplified centrosomes (Fig 4A,B). Similarly, vimentin and mitochondria displacement towards the cell periphery were also supressed following αTAT1 depletion (Fig 4C-F). However, not all membrane-bound organelles displaced in cells with amplified centrosomes were sensitive to the levels of tubulin acetylation. EEA1-positive endosomes and Golgi displacement were not prevented by αTAT1 depletion (Fig EV4C-F), indicating that displacement of these organelles in response to centrosome amplification is regulated by a different mechanism that does not involve tubulin acetylation. Moreover, αTAT1 depletion did not prevent the formation of stable, nocodazole-resistant microtubules in cells with amplified centrosomes (Fig EV4G,H), thus the role of tubulin acetylation in kinesin-1 mediated organelle displacement is independent of increased microtubule stabilization.

**Figure 4.**
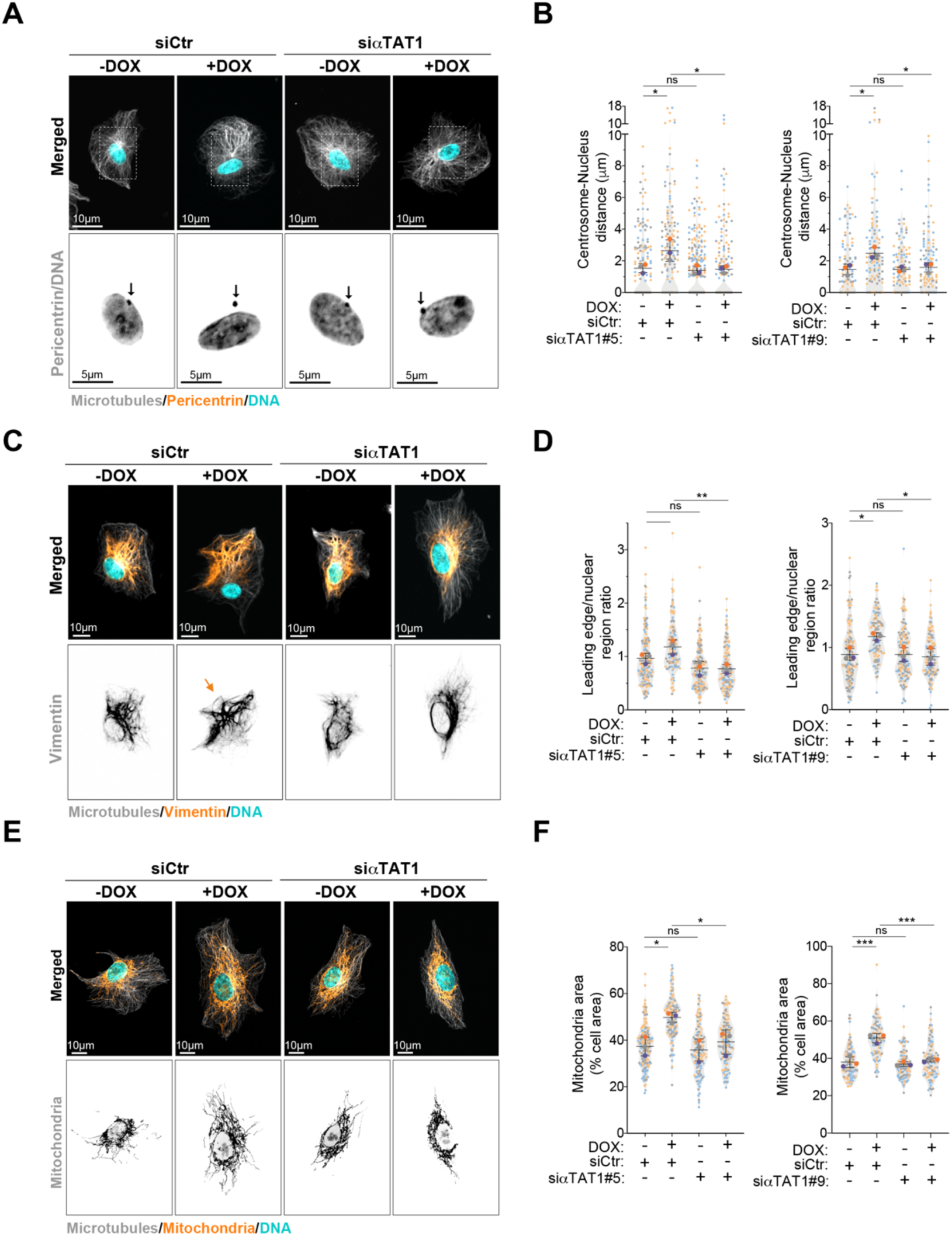
αTAT1-dependent microtubules acetylation controls the displacement of centrosomes, vimentin and mitochondria downstream of centrosome amplification. **A.** Representative images of cells stained for centrosomes (Pericentrin, orange), microtubules (α-tubulin, grey) and DNA (Hoechst, cyan) upon depletion of αTAT1. Black arrows indicate the position of the centrosome(s). Scale bar: 10 µm; inset scale bar: 5 µm. **B.** Quantification of centrosome-nucleus distance (n(-DOX siCtr)=164; n(+DOX siCtr)=158; n(-DOX siαTAT1#5)=235; n(+DOX siαTAT1#5)=173; n(-DOX siCtr)=107; n(+DOX siCtr)=119; n(-DOX siαTAT1#9)=110; n(+DOX siαTAT1#9)=118). **C.** Representative images of cells stained for vimentin (orange), microtubules (α-tubulin, grey) and DNA (Hoechst, cyan) upon depletion of αTAT1. Orange arrows indicate the displacement of vimentin towards cell periphery. Scale bar: 10 µm. **D.** Quantification of vimentin leading edge/nuclear ratio (n(-DOX siCtr)=161; n(+DOX siCtr)=111; n(-DOX siαTAT1#5)=153; n(+DOX siαTAT1#5)=122; n(-DOX siCtr)=144; n(+DOX siCtr)=109; n(-DOX siαTAT1#9)=130; n(+DOX siαTAT1#9)=117). **E.** Representative images of cells stained for mitochondria (MitoTracker, orange), microtubules (α-tubulin, grey) and DNA (Hoechst, cyan) upon depletion of αTAT1. Scale bar: 10 µm. **F.** Quantification of mitochondria area (n(-DOX siCtr)=169; n(+DOX siCtr)=125; n(-DOX siαTAT1#5)=130; n(+DOX siαTAT1#5)=134; n(-DOX siCtr)=169; n(+DOX siCtr)=125; n(-DOX siαTAT1#9)=130; n(+DOX siαTAT1#9)=134). For all graphics error bars represent mean +/- SD from three independent experiments. \**p* < *0.05,* \*\**p* < *0.01, ***p* < *0.001,* ns = not significant (*p* > *0.05*). The following statistic were applied: one- way ANOVA with Tukey’s post hoc test for all graphs.

To test if microtubule acetylation was sufficient to promote intracellular reorganization, we treated cells with H_2_O_2_, which led to similar levels of tubulin acetylation observed in cells with amplified centrosomes but did not change total tubulin levels (Fig 3F,G). H_2_O_2_ treated cells displayed increased centrosome and vimentin displacement (Fig 5A-D), but no effect was observed on endosome displacement or Golgi dispersion, even in cells that exhibited centrosome displacement, further demonstrating that these phenotypes are not co-dependent (Figs 5A and EV5A-C). Because H_2_O_2_ treatment induces mitochondria fragmentation (Fan *et al*, 2010), we were unable to assess mitochondria displacement in this condition (Fig EV5D). Additionally, we tested whether inducing higher levels of tubulin acetylation by treating cells with Tubacin, an inhibitor of the deacetylase HDAC6, or by overexpressing αTAT1 could induce similar phenotypes (Fig EV5E) (Haggarty *et al*, 2003; Shida *et al*., 2010). Indeed, cells treated with tubacin or overexpressing αTAT1 (αTAT1 OE) showed vimentin and mitochondria displacement towards cell periphery (Fig 5E-H). Surprisingly, however, centrosomes remained closely associated with the nuclear envelope, suggesting that high levels of tubulin acetylation alone may not be sufficient to promote their displacement (Figs 5I,J and EV5D). What could explain this difference? We hypothesised that in addition to increased levels, the distribution or orientation of acetylated microtubules could differentially impact centrosome positioning, since as a single organelle it would be more sensitive to the polarisation of pushing forces, where isotropic pushing forces would cancel each other (Fig 6A). To assess this, we divided the cell as rear and front based on the centrosome positioning and quantified the distribution of orientation variations using Fiji (Li *et al*, 2022; Schindelin *et al*, 2012) (Fig 6B,C). Plotting the raw values for the frequency of all orientation variations in the cell front (values >0) and rear (values <0), as well as the normalised orientation values, suggested that the distribution of acetylated microtubules is different between conditions (Fig EV6A,B). Polarisation of orientation variations towards the leading edge can be readily visualised in the rose plots, where a slight increase in polarisation can be observed in cells with amplified centrosomes (+DOX) and those treated with H_2_O_2_. However, this analysis does not take into account the balance of tubulin acetylation between front:rear of the cell. To try to assess this, for each orientation variation, we subtracted the rear frequency from the front frequency (Fig 6D). We found that firstly, it is clear that in all conditions there is an increase in the frequency of orientations towards the front of the cell; and secondly that a clear polarisation of these frequencies towards the leading edge (∼60^0^ to 120^0^) can be observed in cells with amplified centrosomes or treated with H_2_O_2_. Thus, and assuming that increased frequency of tubulin acetylation results in increased kinesin-1 pushing forces, our data could explain why centrosome displacement is only observed in cells with extra centrosomes or treated with H_2_O_2_ since only in these conditions our model predicts directional forces towards leading edge. These results demonstrate that the displacement of centrosomes, vimentin and mitochondria towards the leading edge is regulated by tubulin acetylation and that both levels and distribution of acetylated microtubules could differentially impact these phenotypes.

**Figure 5.**
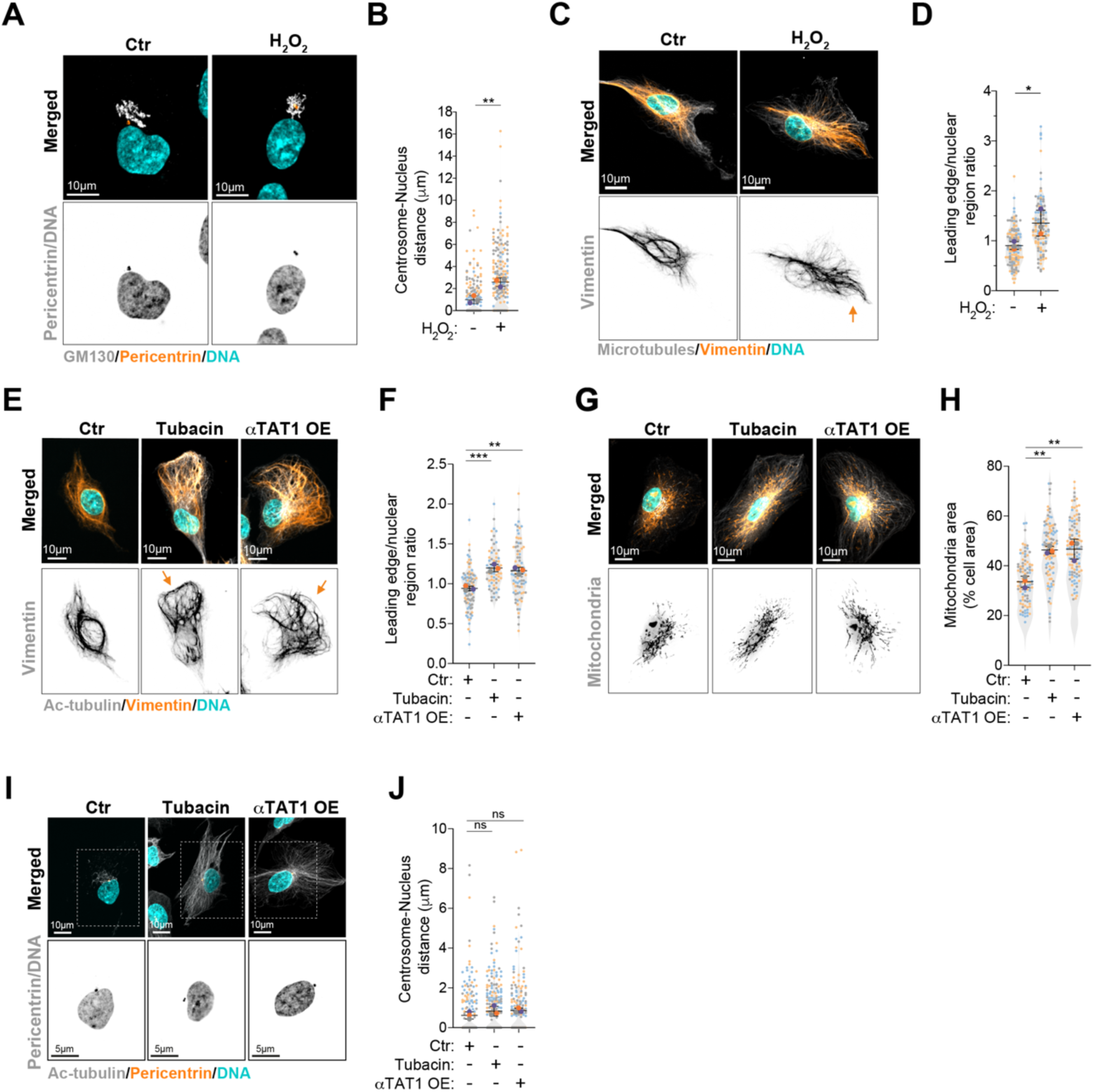
Increased tubulin acetylation is sufficient to promote organelle displacement independently of amplified centrosomes. **A.** Representative images of cells stained for centrosomes (Pericentrin, orange), Golgi (GM130, grey) and DNA (Hoechst, cyan) in cells treated with H2O2 (75µM). Scale bar: 10 µm. **B.** Quantification of centrosome-nucleus distance (n(Ctr)=192; n(H2O2)=211). **C.** Representative images of cells stained for vimentin (orange), microtubules (α-tubulin, grey) and DNA (Hoechst, cyan) treated with H2O2. Orange arrows indicate the displacement of vimentin towards cell periphery. Scale bar: 10 µm. **D.** Quantification of vimentin leading edge/nuclear ratio (n(Ctr)=132; n(H2O2)=117). **E.** Representative images of cells stained for vimentin (orange), acetylated tubulin (Ac-tubulin, grey) and DNA (Hoechst, cyan) in cells treated with Tubacin (5 µM) or overexpression αTAT1-GFP (αTAT1 OE). Orange arrows indicate the displacement of vimentin towards cell periphery. Scale bar: 10 µm. **F.** Quantification of vimentin leading edge/nuclear ratio (n(Ctr)=93; n(Tubacin)=65; n(αTAT1 OE)=85). **G.** Representative images of cells stained for mitochondria (MitoTracker, orange), acetylated tubulin (Ac-tubulin, grey) and DNA (Hoechst, cyan) in cells treated with Tubacin or overexpression αTAT1-GFP (αTAT1 OE). Scale bar: 10 µm. **H.** Quantification of mitochondria area (n(Ctr)=102; n(Tubacin)=106; n(αTAT1 OE)=109). **I.** Representative images of cells stained for centrosomes (Pericentrin, orange), acetylated tubulin (Ac- tubulin, grey) and DNA (Hoechst, cyan) in cells treated with Tubacin or overexpression αTAT1-GFP (αTAT1 OE). Scale bar: 10 µm. **J.** Quantification of centrosome-nucleus distance (n(Ctr)=215; n(Tubacin)=260; n(αTAT1 OE)=212). For all graphics error bars represent mean +/- SD from three independent experiments. \**p* < *0.05,* \*\**p* < *0.01, ***p* < *0.001,* ns = not significant (*p* > *0.05*). The following statistic were applied: unpaired *t* test for graphs in B and D and one-way ANOVA with Tukey’s post hoc test for graphs in F, H and J.

**Figure 6.**
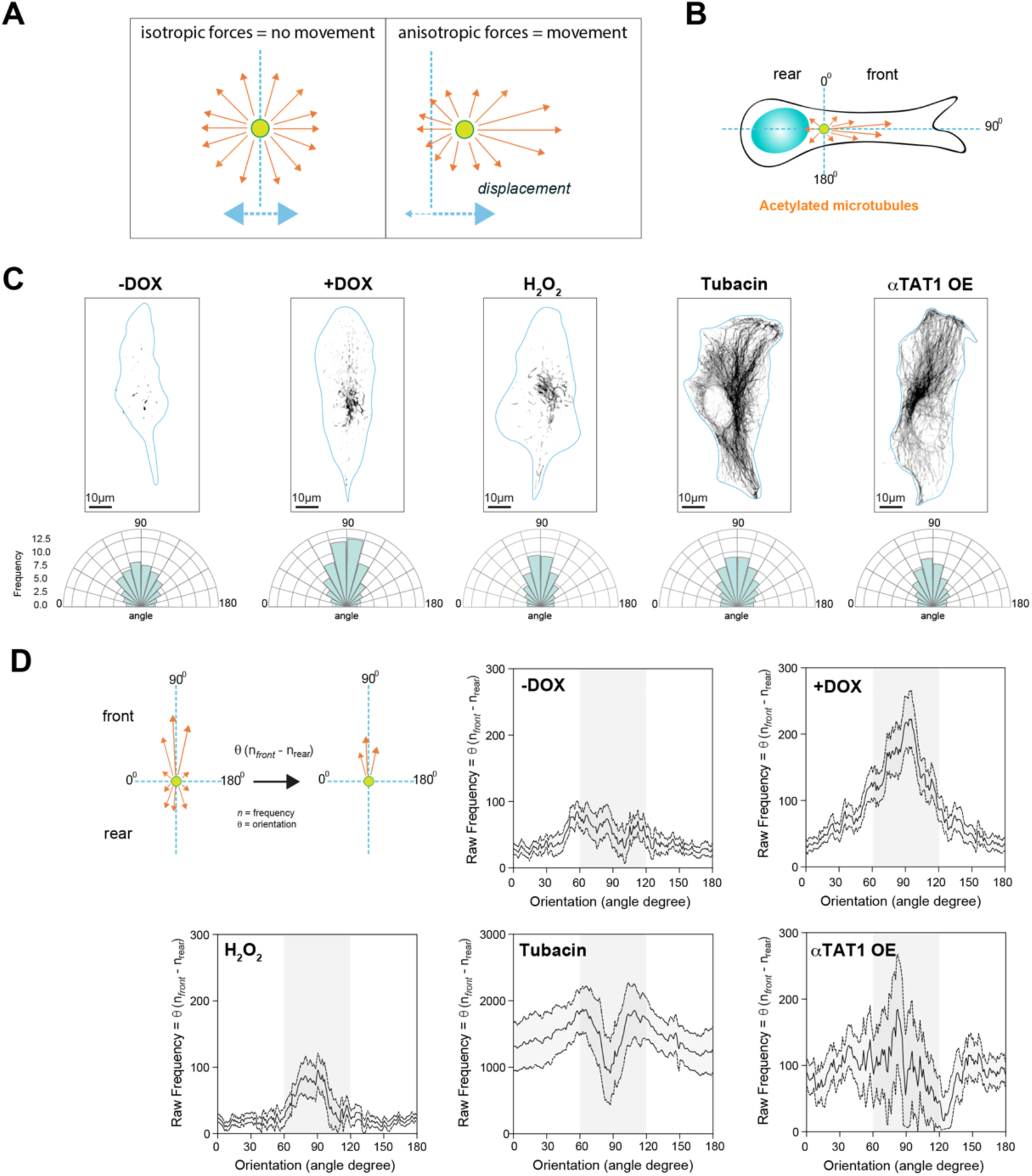
Polarization of acetylated microtubules in cells varies with different acetylated tubulin-inducing conditions. **A.** Scheme representing how distribution of forces could impact movement/displacement of single organelles (e.g. centrosome). **B.** Scheme depicting how orientation of acetylated microtubules was determined in cells. **C.** Top: Representative images of acetylated microtubules (grey) in control cells (-DOX), cells with amplified centrosomes (+DOX), treated with H2O2 or Tubacin and overexpressing αTAT1-GFP (αTAT1 OE). Scale bar: 10µm. Bottom: Rose plots displaying the frequency of acetylated microtubule orientation in the cell front. **D.** Quantification of the distribution of acetylated microtubule orientation upon subtracting rear values from front values (n(-DOX)=61; n(+DOX)=48; n(H2O2)=83; n(Tubacin)=39; n(αTAT1 OE)=63).

### Intracellular reorganization in cells with amplified centrosomes correlates with enhanced nuclear deformability

Nucleus-associated vimentin confers a protective role to the nucleus against mechanical stress and its loss enhances nuclear deformability (Patteson *et al*, 2019a; Patteson *et al*, 2019b). Thus, we hypothesized that intracellular reorganization resulting in vimentin displacement towards the cell periphery could promote nuclear deformability. Nucleus aspect ratio was used as proxy for deformability, where 1=perfect circle (Fig 7A). When plated in 3D confined collagen-I matrices, cells with amplified centrosomes displayed lower nuclear circularity compared to control cells, suggesting increased nuclear deformability (Fig 7B,C). Although actin has long been proposed to play a role in mediating nucleus deformation in different cells types (Thiam *et al*, 2016), treatment with latrunculin-A did not prevent nuclear deformability in cells with amplified centrosomes. By contrast, microtubule depolymerization with nocodazole prevented nuclear deformability (Fig EV7A,B). Moreover, KIF5B depletion also blocked increased nuclear deformability (Fig 7B,C). Similarly to what was observed for organelle displacement, p150^glued^ depletion did not impact cells with extra centrosomes but was sufficient to increase nuclear deformability in control cells (Fig 7B,C). These results support a model where changes in the balance of microtubule motor protein activity driven by centrosome amplification enhances nuclear deformability, likely as a result of vimentin displacement towards the cell leading edge (Fig 2H).

**Figure 7.**
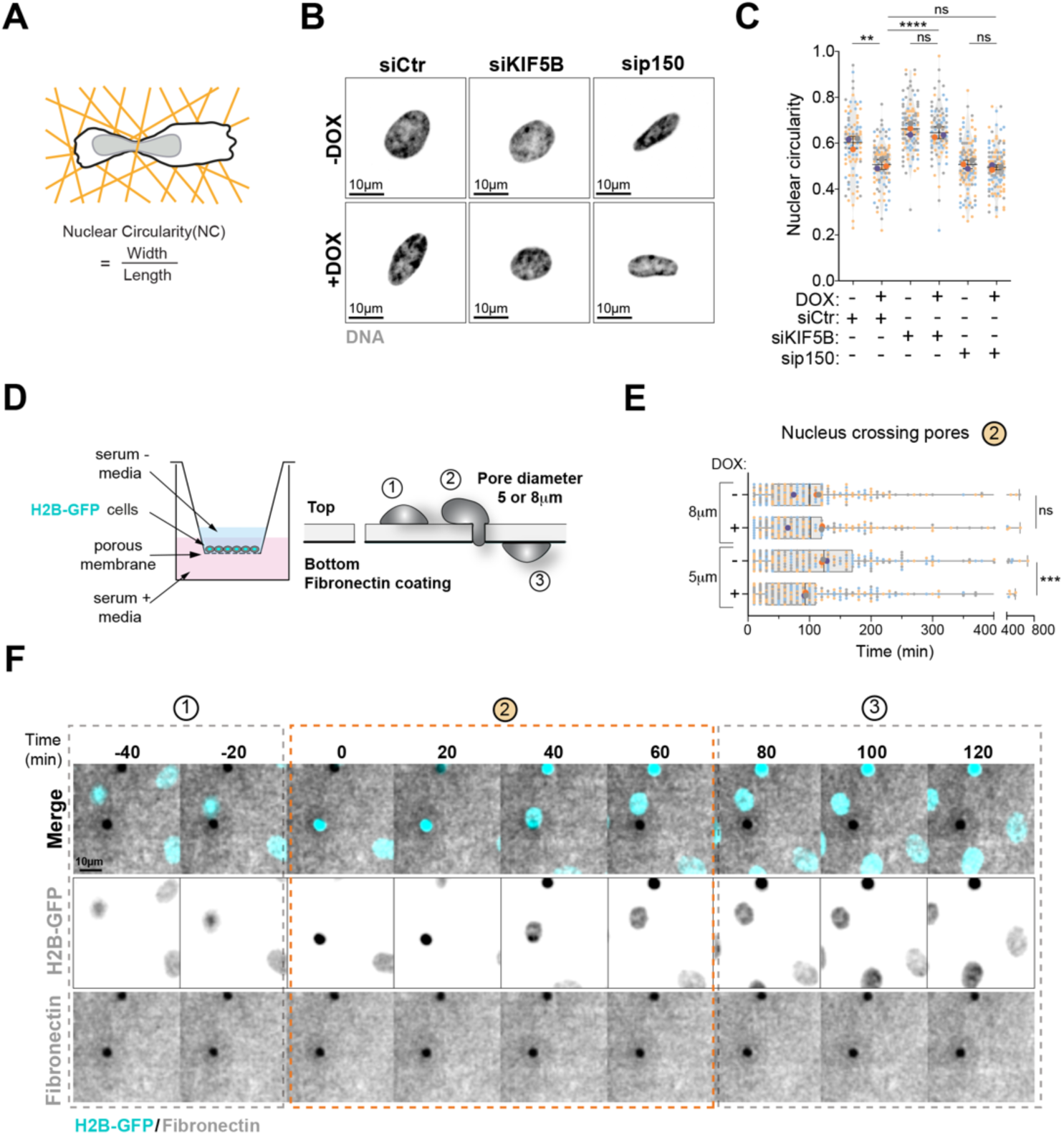
Centrosome amplification enhances nucleus deformation which leads to efficient nucleus translocation through restrictive constrictions. **A.** Representative scheme of nucleus aspect ratio quantification. **B.** Representative images of the nucleus (Hoechst, grey) in cells upon depletion of KIF5B and p150. Scale bar: 10µm. **C.** Quantification of nucleus aspect ratio (n(-DOX siCtr)=95; n(+DOX siCtr)=107; n(-DOX siKIF5B)=111; n(+DOX siKIF5B)=91; n(-DOX sip150)=115; n(+DOX sip150)=117). **D.** Representative scheme of the Transwell system and nucleus crossing constrictions. **E.** Quantification of time spent by nucleus to cross 5 µm- or 8 µm- diameter constrictions (-DOX(5µm)=228; +DOX(5µm)=259; -DOX(8µm)=231; +DOX(8µm)=215). **F.** Still images from live cell imaging depicting cell nucleus (H2B-GFP, cyan) crossing pores on membranes coated with fibronectin (grey). Time spent in phase 2 is depicted in the graph in E. Scale bar: 10 µm. For all graphics error bars represent mean +/- SD from three independent experiments. \*\**p* < *0.01, ***p* < *0.001, ****p* < *0.0001,* ns = not significant (*p* > *0.05*). The following statistic were applied: unpaired *t* test for graph in E (analyses of migration in 8 µm and 5 µm was done separately) and one-way ANOVA with Tukey’s post hoc test for graph in C.

During cell migration through confined spaces, the nucleus, which is the largest and stiffest cellular organelle, constitutes a burden for cells (Denais *et al*, 2016; Raab *et al*, 2016). Thus, we hypothesized that the increased nuclear deformability in cells with amplified centrosomes could facilitate migration through confined spaces. To test this, we utilized a Transwell assay in which cells were seeded onto a membrane and allowed to migrate through pores of different sizes. Using RPE-1.iPLK4 cells expressing H2B-GFP to visualize the nucleus, the speed of nuclear translocation through these pores was assessed by live-cell imaging (Fig 7D). We found that the time for the nucleus to cross the larger 8µm pores was similar in cells with normal and amplified centrosomes (-DOX=97.01 ± 5.779min; +DOX= 97.07 ± 6.41min). However, cells with amplified centrosomes migrated faster through smaller 5µm pores (- DOX=123.4 ± 7.345min; +DOX= 92.74 ± 5.77min), suggesting that increased nuclear deformability provides an advantage in constrained environments (Fig 7E,F). Altogether, these results suggest that changes in organelle organization in cells with extra centrosomes, could enhance nuclear deformation to facilitate migration through confined spaces.

## Discussion

Here, we demonstrate that centrosome amplification is sufficient to change intracellular organisation, a process that requires kinesin-1 mediated transport and partly regulated by increased tubulin acetylation. This intracellular reorganisation in cells with amplified centrosomes increases nuclear deformability and facilitates nuclear migration through small constrictions. The differential impact of acetylated tubulin on organelle distribution highlights more complex sensing and response mechanisms by which organelles read the tubulin code.

The centrosome positioning at the cell center and in close proximity to the nucleus has long been proposed to result from an equilibrium of pulling and pushing forces exerted by microtubule motors (Bornens, 2008). In late G2, Inhibition of dynein leads to centrosome displacement away from the nucleus in a kinesin-1-dependent manner, from the cell center towards cell periphery (Splinter *et al*., 2010), demonstrating that dynein functions as a brake that counteracts kinesin-1 forces. Interestingly, we found that induction of centrosome amplification is sufficient to drive the displacement of centrosomes towards cell periphery, phenocopying what has been observed in cells upon dynein inhibition. Depletion of the kinesin-1 KIF5B prevents displacement of the centrosomes, implying that pushing forces on centrosomes are mediated by KIF5B and that these forces overcome the pulling activity of dynein. Supporting this idea, disconnecting centrosomal microtubules from the nuclear envelope by disrupting the LINC complex further exacerbates centrosome displacement in cells with amplified centrosomes. We also observed that, in response to centrosome amplification, several intracellular compartments are displaced towards the cell periphery, namely endosomes, mitochondria, vimentin and Golgi. This global reorganization is consistent with previous observations in pancreatic cancer cells showing that centrosome amplification leads to the dispersion of late endosomes/multivesicular bodies towards the cell periphery (Adams *et al*., 2021). Displacement of endosomes and vimentin have also been shown to require kinesin-1 (Gyoeva & Gelfand, 1991; Liao & Gundersen, 1998; Nath *et al*, 2007; Schmidt *et al*, 2009), potentially highlighting a kinesin-1 mediated global reorganization of the cytoplasm in cells with amplified centrosomes.

Systematic analyses of different intracellular compartments revealed that reduction of acetylated tubulin levels, via depletion of αTAT1, prevented the displacement of centrosomes, vimentin and mitochondria towards cell periphery, indicating that enhanced tubulin acetylation plays a role in the relocation of these intracellular compartments. By contrast, depletion of αTAT1 had no significant impact on endosomes and Golgi reorganization, suggesting that other microtubule PTMs and/or adaptor proteins, which link organelles to microtubules, could specifically affect relocation of these organelles in cells with amplified centrosomes (Akhmanova & Hammer, 2010; Barlan & Gelfand, 2017; Cross & Dodding, 2019). It is also possible that other kinesin motors, such as kinesin-3, could transport endosomes independently of tubulin acetylation (Bielska *et al*, 2014; Wedlich-Soldner *et al*, 2002). Exactly how tubulin acetylation, which occurs in the microtubule lumen, favors kinesin-1 mediated organelle transport remains unclear. Our data demonstrates that microtubule stabilization is unlikely to be the answer since αTAT1 depletion in cells with amplified centrosomes does not affect the number of nocodazole-resistant microtubules. Therefore, microtubule stabilization is unlikely to play a role in kinesin-1 mediated displacement of centrosomes, vimentin and mitochondria.

Unexpectedly, we found that not only the levels but also the distribution of acetylated microtubules could contribute to organelle displacement. High levels of tubulin acetylation induced by Tubacin or αTAT1 overexpression induced the displacement of vimentin and mitochondria towards cell periphery, but not centrosomes. This contrasts with what was observed in cells with amplified centrosomes or treated with H_2_O_2_, which induce lower levels of tubulin acetylation. We propose that an anisotropic distribution of forces is required to displace centrosomes. Indeed, we observed a polarized distribution of acetylated microtubules towards the leading edge in cells harboring amplified centrosomes or treated with H_2_O_2_, suggestive of anisotropic distribution of kinesin-1-mediated pushing forces required for centrosome displacement. While our data is only suggestive of such model, it raises an important issue when assessing the role of microtubule PTMs in cells, that not all conditions that increase specific PTMs may elicit the exact same phenotype and that distribution of modified microtubules should be taken into consideration.

What are the consequences of this intracellular reorganization in cells with amplified centrosomes? Extensive changes in cell shape occur as cells migrate, and this is accompanied by relocations of several organelles and cellular compartments (Bornens, 2008). It is plausible that during migration, displacement of vimentin towards cell periphery could lead to nuclear deformation to facilitate migration through confined spaces. Consistently, cells with amplified centrosomes display increased nuclear deformability in confined 3D collagen gels and move faster through smaller pores. Extreme nuclear deformability has also been observed in invasive MCF10A cells with extra centrosomes migrating through thin invasive protrusions (Godinho *et al*., 2014). Thus, it is tantalizing to propose that vimentin displacement, rather than loss, could help migration through confined spaces in a more controlled manner while preventing extensive nuclear rupture and DNA damage observed in vimentin knock-out cells (Patteson *et al*., 2019b).

It has been recently proposed that binding of the ER to glutamylated microtubules plays a role in orchestrating the movement and positioning of several organelles, in an attempt to centralize intracellular organization (Zheng *et al*., 2022). However, this is unlikely to be a general feature. While ER tubules were shown to slide preferentially along acetylated microtubules (Friedman *et al*., 2010), our findings demonstrate that not all organelles respond to tubulin acetylation. This indicates that, depending on the context, individual organelles must have their own sensing and response mechanisms to ensure fine-tuning of their distribution in cells. We propose that this fine-tuning enables cells to adapt to different stimuli and environments.

## Acknowledgements

We are grateful to all the members of the Godinho lab for comments and discussion of the manuscript. We thank Edgar Gomes for providing the KASH-DL and KASH2 constructs. P.M. and S.S.W. were supported by Medical Research Council Grants (MRC, MR/M010414/1 and MR/T000538/1). Bongwhan Yeon was supported by a Cancer Research UK (CRUK) PhD studentship. S.A.G. is a fellow of the Lister Institute and is supported by the MRC (MR/T000538/1). This work was supported by a Cancer Research UK Centre Grant to Barts Cancer Institute (C355/A25137).

The authors declare that there is no conflict of interest.

## Materials and Methods

### Cell culture

RPE-1 (human retinal epithelial) cells were grown in Dulbecco’s modified Eagle’s medium/nutrient mixture F-12 Ham (DMEM-F12; Sigma) supplemented with 10% Fetal Bovine Serum (FBS; Gibco) and 100 U/ml Penicillin/Streptomycin (P/S; Gibco) and maintained at 37°C with 5% CO_2_ atmosphere. Tetracycline-free FBS (Gibco) was used to grow cells expressing the PLK4 tet-inducible construct. The FBS was heat inactivated at 56°C water bath for 30 min.

### Plasmids and cell lines

RPE-1.iPLK4 and RPE-1.iPLK4^1-608^ cell lines were generated using pLenti-CMV-TetR-Blast lentiviral vector (Addgene, 17492) and selected using Blasticidin (10 µg/mL). Post-selection, cells were then infected with a lentiviral vector containing either PLK4 WT or PLK4^1-608^ mutant cDNA which had been previously cloned into the pLenti-CMV/TO-Neo-Dest vector and selected using Geneticin (200 µg/mL)(Godinho *et al*., 2014). Cells expressing the PLK4 WT and PLK4^1-608^ mutant transgenes were then induced for 48 h using 2 µg/mL of Doxycycline. The LV-GFP plasmid (Addgene, 25999) was used to express H2B-GFP and cells were selected by FACS (Beronja et al., 2010). eGFP-αTAT1 was prepared from pEF5B-FRT-GFP-αTAT1 (Addgene, 27099) by PCR with BamHI and SalI restriction sites on the 5’ and 3’ end respectively and cloned into pLV-eGFP (addgene, 36083).

### Lentiviral generation

To generate lentivirus, HEK-293 cells were plated in antibiotic free medium. Transfection of the appropriate lentiviral plasmid in combination with Gag-Pol (psPAX2, Addgene, 12260) and VSV-G (VSV-G: pMD2.G, Addgene, 12259) was performed using Lipofectamine 2000® (Thermo Fisher Scientific), as per the manufacturer’s specifications. The resultant lentivirus was harvested 24 h and 48 h post infection, passed through a 0.4 µM syringe filter and stored in cryovials at −80°C. For infection, the appropriate lentivirus was then mixed with 8 μg/mL polybrene before being added to the cells in a dropwise fashion. Infection was repeated the following day and antibiotic selection started 24 h after final infection.

### Chemicals

Chemicals and treatments were performed as follows: 2 µg/mL Doxycycline hyclate (DOX; Sigma) treatment for 48 h, 75 µM hydrogen peroxide (H_2_O_2_; Sigma) treatment for 4 h, 0.5 mM of Apocynin (Santa Cruz) treatment for 72 h added at the same time as DOX, 0.1 µM LatrunculinA (LatA; Sigma) treatment for 5 h, 10 µM Nocodazole (Noc; Sigma) treatment for 5 h to completely depolymerize microtubules and 2 µM Nocodazole for 30 min to assess the numbers of microtubules resistant to Noc, 5 µM of Tubacin (Sigma) was added for 4 h before fixation.

### siRNA transfection

siRNA transfection was performed in antibiotic free growth medium using Lipofectamine® RNAiMAX (Thermo Fisher) as per the manufacturer’s instructions. Briefly, cells were grown in a 6-well plate until reaching ∼60% confluency. Prior to transfection, growth medium was replaced with 2 mL of fresh growth medium without antibiotics. For each well, the transfection solution was prepared as followed: 10 µL of Lipofectamine® RNAi MAX Transfection Reagent was diluted in 250 µL of Reduced Serum Medium Opti-MEM® (Thermo Fisher) in a sterile 1.5 mL microcentrifuge tube and 5 µL of siRNA at 20 µM was diluted in 250 µL of Opti-MEM® in a separate sterile 1.5 mL microcentrifuge tube. Tubes were then incubated at room temperature (RT) for 5 min for equilibration. Opti-MEM® solution containing siRNA was then added dropwise onto the tube containing the lipofectamine RNAi MAX solution and incubated for 20 min at RT to allow liposome formation. The solution was then added dropwise onto the 6-well and incubated for 6 h. After 6 h media was refreshed, and cells were analyzed 72h post-transfection. siRNAs used in this study are listed below:

| siRNA | Reference | Company |
| --- | --- | --- |
| siControl (Ctr) | 1027310 | Qiagen |
| siKIF5B SmartPool | L-008867-00-0005 | Dharmacon |
| siP150glued SmartPool | L-012874-00-0005 | Dharmacon |
| siαTAT1 #5 | SI03124660 | Qiagen |
| siαTAT1 #9 | SI04145162 | Qiagen |

### RNA extraction, quantification, and cDNA generation

Total RNA extraction was carried out using RNeasy kit (Qiagen) according to the manufacturer’s instructions. Eluted RNA was then stored at −80°C. RNA concentration was determined using Nanodrop 1000 spectrophometer (Thermo Fisher, USA). For cDNA generation, 500 ng of total RNA was mixed with 2 µL of random primer mix (New England Biolabs, UK) in RNase free PCR strips (Thermo Fisher, USA). RNase-free water was added to a final volume of 16 µL. The mixture was heated for 3 min at 65°C in a PCR machine. Tubes were then placed on ice immediately for few minutes. After that, 2 µL of RT buffer, 1 µL of RNase inhibitor (New England Biolabs) and 1 µL of reverse transcriptase was added on top of 16 µL of the extracted RNA. Tubes were incubated at 42°C for 1 h for reverse transcriptase elongation and at 90°C for 15 min for reverse transcriptase inactivation to create a pool of cDNA. cDNA was then stored at −20°C.

### qRT-PCR

A PCR cocktail was generated by adding 9 µL of nuclease-free water (Thermo Fisher), 30 µL of 2X Power SYBR® Green PCR Master Mix (Thermo Fisher) and 3 µL of gene specific forward and reverse primers at 10 µM to generate a final volume of 45 µL. cDNA was diluted by adding 4.5 µL of nuclease-free water onto 0.5 µL of cDNA to make a final volume of 5 µL. 15 µL of the PCR cocktail was added onto each well in triplicate in a 96-well plate and 5 µL of the diluted cDNA was then added to all wells in triplicate, giving a final volume of 20 µL in each well. The 96-well plate was then sealed and centrifuged for few seconds to spin down the mixture. The Ct values acquired from the qRT-PCR reaction were analyzed by using comparative Ct method (2-ΔΔCt) and GAPDH was used as a housekeeping gene for normalization.

### Indirect immunofluorescence

1-1.5 x 10^4^ cells were seeded in a final volume of 50-80 µL on an 18 mm-diameter glass coverslip in a 12-well plate. The plate was then incubated at 37°C for 30 min to allow cell attachment. Once cells attached, 1 mL of the growth medium was added and the plate was incubated at 37°C overnight. On the following day, growth medium was aspirated and coverslips were washed with PBS once and fixed immediately with either 4% PFA+PBS at RT for 15 min or with 99.9% ice-cold methanol at −20°C for 10 min. After fixation, all steps were carried out at RT. Cells were incubated with permeabilization buffer (PBS +0.2% Triton X-100) for 5 min. After permeabilization, cells were blocked with 1 mL of blocking buffer (PBS, 5% BSA, and 0.1% Triton X-100) for 30 min. 30 µL of the diluted primary antibodies were added onto the coverslip and incubated for 30 min, after which another 30 µL was added for another 30 min to avoid coverslips to dry. Next, coverslips are washed twice with PBS and secondary antibodies (Alexa Fluor conjugated; Molecular Probes) incubation was performed in the same way as the primary antibodies in the dark. Hoechst 33342 solution was used at 1:10000 dilution to stain DNA in the dark. Coverslips were then mounted on a drop of ProLong Gold antifade reagent on a microscope slide. Primary and secondary antibodies and molecular probes used in this paper are listed below:

| Antibody (Clone) | Species | Catalogue number | Manufacturer | Dilution | Fixation |
| --- | --- | --- | --- | --- | --- |
| Vimentin | Rabbit | 5741 | Cell signalling | 1:200 | Methanol (MeOH) |
| EEA1 | Rabbit | 2411S | Cell signalling | 1:100 | Paraformaldehyde (PFA) |
| Pericentrin | Rabbit | Ab4448 | Abcam | 1:1500 | MeOH |
| α-tubulin (DM1α) | Mouse | T9026 | Sigma Aldrich | 1:1000 | MeOH |
| α-tubulin FITC-conjugated | Mouse | F2168 | Sigma Aldrich | 1:200 | MeOH |
| Acetylated tubulin (6-11B-1) | Mouse | T6793 | Sigma Aldrich | 1:2500 | MeOH |
| GM130 | Mouse | 610822 | BD biosciences | 1:100 | MeOH/PFA |
| EB1 | Mouse | 610535 | BD biosciences | 1:500 | MeOH/PFA |
| Nesprin-2 | Mouse | sc-365097 | Santa Cruz | 1:200 | PFA |
| Nesprin-3 | Rabbit |  | Gift from A. Sonnenberg | 1:200 | PFA |
| Anti-Mouse Alexa Fluor 488 | Goat | A11001 | Life Technologies | 1:1000 |  |
| Anti-Mouse Alexa Fluor 647 | Goat | A21235 | Thermofisher Scientific | 1:1000 |  |
| Anti-Rabbit Alexa Fluor 488 | Goat | A11008 | Life Technologies | 1:1000 |  |
| Anti-Mouse Alexa Fluor 568 | Goat | A11004 | Life Technologies | 1:1000 |  |

| Probe/dye | Catalogue number | Manufacturer | Dilution | Fixation |
| --- | --- | --- | --- | --- |
| Alexa Fluor 568 Phalloidin | A12380 | Thermo Fisher | 1:500 | PFA |
| Mitotracker | M7510 | Thermo fisher | 1:10000 | PFA |

### Western blotting

Cells were collected and resuspended in 100 µl of RIPA buffer (Thermo Fisher Scientific) with added protease inhibitors (Roche; 1 tablet/10 ml RIPA). Protein concentration was quantified using the Bio-Rad DC protein assay and 15 µg of protein was loaded per well. Protein samples were resuspended in Laemmli buffer and separated on SDS-PAGE and transferred onto PVDF membranes. Western blots were developed using SRX-101A Konica Minolta and scanned. Antibodies used for western blot analysis are listed below.

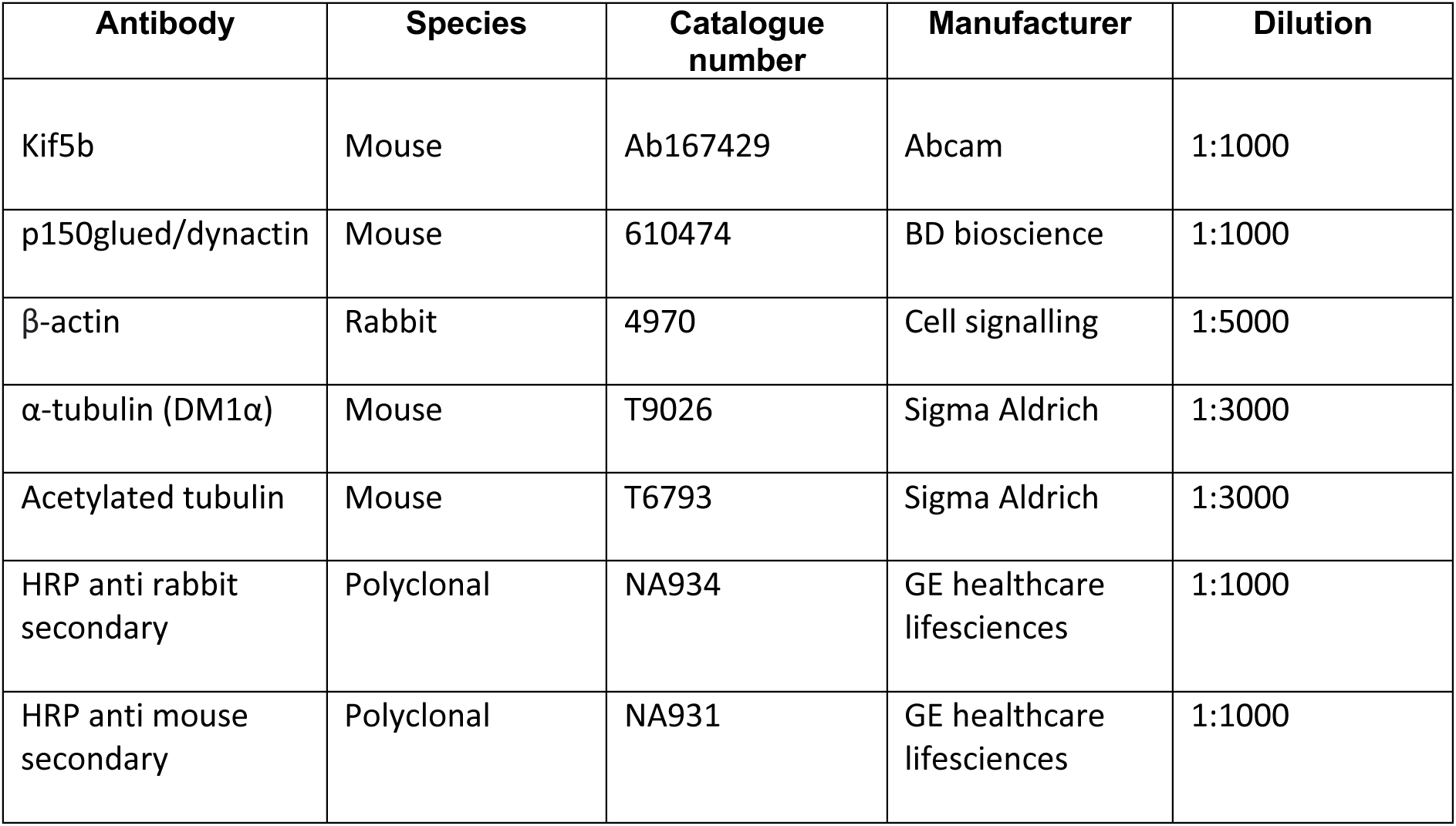

### 3D Collagen gels

Collagen gels were performed as previously described (Infante *et al*, 2018). Briefly, glass coverslips were layered with 15 µL of a 2.2 mg/mL type-I collagen solution (bottom layer). Polymerization was induced at 37 °C for 3 min. Then, a cell suspension (1.5–2.5 × 10^5^ cells/mL) was added to the bottom layer and cultures were incubated for 30 min at 37 °C to allow cells to adhere to the collagen gels. Growth medium was gently removed and a 2.2 mg/mL type-I collagen solution was polymerized on top of the cells (top layer). After polymerization at 37 °C for 90 min, growth medium was added to the cultures. Z-stacks of images were acquired with an inverted Nikon microscope coupled with a spinning disk confocal head (Andor) with a 60x objective.

### Quantifications of indirect immunofluorescence images

For centrosome number quantification, cells were stained with DNA and centrin2 and number of centrosomes were quantified in mitosis: 4 centrioles = 2 centrosomes (normal) and >5 centrioles = amplified centrosomes. Centrosome-nucleus distance was manually assessed using ImageJ by drawing a line between nucleus edge and centrosome(s) center. For nucleus aspect ratio quantification, cells were stained with a DNA dye (Hoechst) and ImageJ was used to assess the width and height of the nucleus (ratio=1, round nucleus). To quantify endosomes-nucleus distance, cells were stained with EEA1 and DNA. Individual endosome distance to the nucleus was manually determined using ImageJ ‘find maxima plugin’ to determine endosomes coordinates. Mean endosomes distance per cell was then calculated as the average of all endosomes for each cell. For the quantification of vimentin displacement, cells were stained for vimentin and microtubules and vimentin fluorescence intensity was determined as previously described (Leduc & Etienne-Manneville, 2017). Briefly, vimentin fluorescence intensity was calculated as the ratio between fluorescence intensity at the leading edge (15 µm from the leading edge) and in a 20µm-radius perinuclear region (ratio<1= vimentin associated with the nucleus; ratio>1= vimentin displaced towards cell leading edge). For Golgi area quantification, cells were stained with GM130 and DNA. Golgi area was manually determined using ImageJ ‘freehand draw’ and ‘measure’ options. For mitochondria area quantification, cells were stained with MitoTracker (in order to visualize mitochondria) and with phalloidin (in order to visualize F-actin and determine cell area and border). Mitochondria spreading was determined as the ratio between mitochondria area over total cell area. For all imaging experiments, cells were imaged with an Eclipse Ti-E inverted microscope (Nikon) equipped with a CSU-X1 Zyla 4.2 camera (Ti-E, Zyla; Andor), including a Yokogawa Spinning Disk, a precision motorized stage, and Nikon Perfect Focus, all controlled by NIS-Elements Software (Nikon). 60× 1.45-NA oil objective was used to acquire images.

### Acetylated microtubule orientation variations

Images for -DOX, +DOX, H2O2 and αTAT1 overexpression were collected using an Eclipse Ti-E inverted microscope (Nikon) equipped with a CSU-X1 Zyla 4.2 camera (Ti-E, Zyla; Andor) and tubacin images were collected using an LSM 880 point-scanning confocal microscope (Zeiss). Images of individual cells were transformed in Fiji to have all cells aligned horizontally, with the leading edge on the right and cell rear on the left. Using the acetylated tubulin signal as a mask, the orientations of acetylated microtubules was calculated using the OrientationJ plugin for Fiji (Puspoki *et al*, 2016; Rezakhaniha *et al*, 2012) for the front of the cell (as defined by the centrosome to the leading edge) and the cell rear (defined as the centrosome to the rear of the cell). The front and rear frequencies were summed to calculate the total frequency of all orientations and used to normalize the data for each cell. The difference between front and rear orientation frequency was calculated by subtracting the rear frequency from the front frequency for each orientation.

### Reactive oxygen species (ROS) quantification by live-cell imaging

To measure ROS levels in live cells, 4 x 10^4^ cells (-DOX and H_2_O_2_) and 5 x 10^4^ cells (+DOX and +DOX +Apocynin) were seeded overnight in 8-well glass bottom chambers (iBidi). On the following day, cells were washed with 1x PBS twice and incubated for 20 min in dark at 37°C with 20 µM of H_2_DCFDA (2’,7’- dichlorodihydrofluorescein diacetate; I36007, Thermo Fisher) diluted in serum-free medium. After incubation with H_2_DCFDA, cells were incubated for 5 min in the dark at 37°C with Hoechst 33342 diluted 1:10000 in full growth medium. Wells were then washed with 1x PBS twice and 300 µl of growth medium was added per well. Cells stained with carboxy-H_2_DCFDA were immediately imaged on an Eclipse Ti-E inverted microscope (Nikon) equipped with a CSU-X1 Zyla 4.2 camera (Ti-E, Zyla; Andor), including a Yokogawa Spinning Disk, a precision motorized stage, and Nikon Perfect Focus, all controlled by NIS-Elements Software (Nikon). The microscope was enclosed within temperature- and CO_2_-controlled environments that maintained an atmosphere of 37°C and 5% humidified CO_2_ for live-cell imaging. 60× 1.45-NA oil objective was used to capture images at multiple fields (∼15 fields) z-stack images were captured with 0.5 μm step size and the step size was calculated to minimal pixel overlapping between steps. This procedure was repeated for each condition. ‘nd’ files containing z-stack images were directly opened in the Fiji software. SUM projection was applied to obtain a 2D image and fluorescence intensity was quantified per cell per field. Raw integrated density of multiple cells was measured. To obtain mean total fluorescence intensity per cell in a field, the total fluorescence intensity was divided by the total number of cells per field. 5-10 fields were analyzed to have a total number of ∼30 cells per condition for each experiment.

### Quantification of acetylated tubulin

‘nd’ files containing z-stack images were directly opened in the Fiji software. SUM projection was applied to obtain a 2D image. To quantify total fluorescent intensity of single cells, the boundaries of single cells within an image were outlined using the ‘freehand’ selection tool in the Fiji software. By using the ‘measure’ command, raw integrated density, mean and area of a single cell were measured. After that, a region without fluorescence outside the cell (background) was outlined and measured to obtain mean background fluorescence. Background-corrected total fluorescence intensity of a single cell was determined using the formula = Raw integrated density - (Area of selected cells x Mean fluorescence of background reading).

### Quantification of nocodazole resistant microtubules

To quantify the number of microtubules that resist nocodazole treatment we followed a previously published protocol(Xu *et al*., 2017). Briefly, cells were plated on glass coverslips overnight. The following day cells were treated with 2 μM of nocodazole in growth medium at 37°C for 30 min. Coverslips were then washed in the extraction buffer (60 mM PIPES, 25 mM HEPES, 2 mM MgCl2, 10 mM EGTA, pH 7.0) by rinsing quickly. To extract soluble tubulin, coverslips were immersed in the same extraction buffer containing 0.2% Triton X-100 and 2 μM of nocodazole for 1 min at room temperature. Cells were quickly fixed in cold methanol at −20°C for 10 min. Next, normal immunofluorescence protocol to stained for microtubules, acetylated tubulin and DNA was applied. Cells were imaged using an inverted Zeiss LS880 confocal and a 60x objective. Number of microtubules were quantified manually in Fiji.

### Transwell migration assay

RPE-1 cells stably expressing H2B-GFP were grown on transwell chambers (iBidi). Briefly, the bottom of the upper chamber is a cell-permeable membrane with 5µm or 8µm diameter pore size holes allowing cells to migrate through the chamber. Cell-permeable membrane was coated on their external side, where cells attach, with 20µg/mL fibronectin and 10µg/mL fluorescent conjugated fibronectin solution. Cells were cultured in the upper chamber in serum-free medium while serum-containing medium was added to the wells that function as an attractant to cells, allowing efficient cell migration through the pores. Transwells were imaged for 12-16 hr on an Eclipse Ti-E inverted microscope (Nikon) equipped with a CSU-X1 Zyla 4.2 camera (Ti-E, Zyla; Andor), including a Yokogawa Spinning Disk, a precision motorized stage, and Nikon Perfect Focus, all controlled by NIS-Elements Software (Nikon). The microscope was enclosed within temperature- and CO_2_-controlled environments that maintained an atmosphere of 37°C and 5% humidified CO_2_ for live-cell imaging. Movies were acquired with a Plan Apochromat 60× 1.45-NA oil objective with a 0.13-mm working distance.

### Statistical Analysis

Graphs and statistics were generated using Prism 8 (GraphPad Software) where results are presented as mean ± standard deviation (SD) unless otherwise stated. Statistical analysis was performed using one-way ANOVA with a Tukey’s post hoc test, paired *t* test or one sample *t* test for normalized data (using a hypothetical mean of 1). Different tests utilized are highlighted on the figure legends. Significance is equal to *p<0.05, **p<0.01, ***p<0.001 and ****p<0.0001.

## Supplementary Figures + Legends

**Figure EV1.**
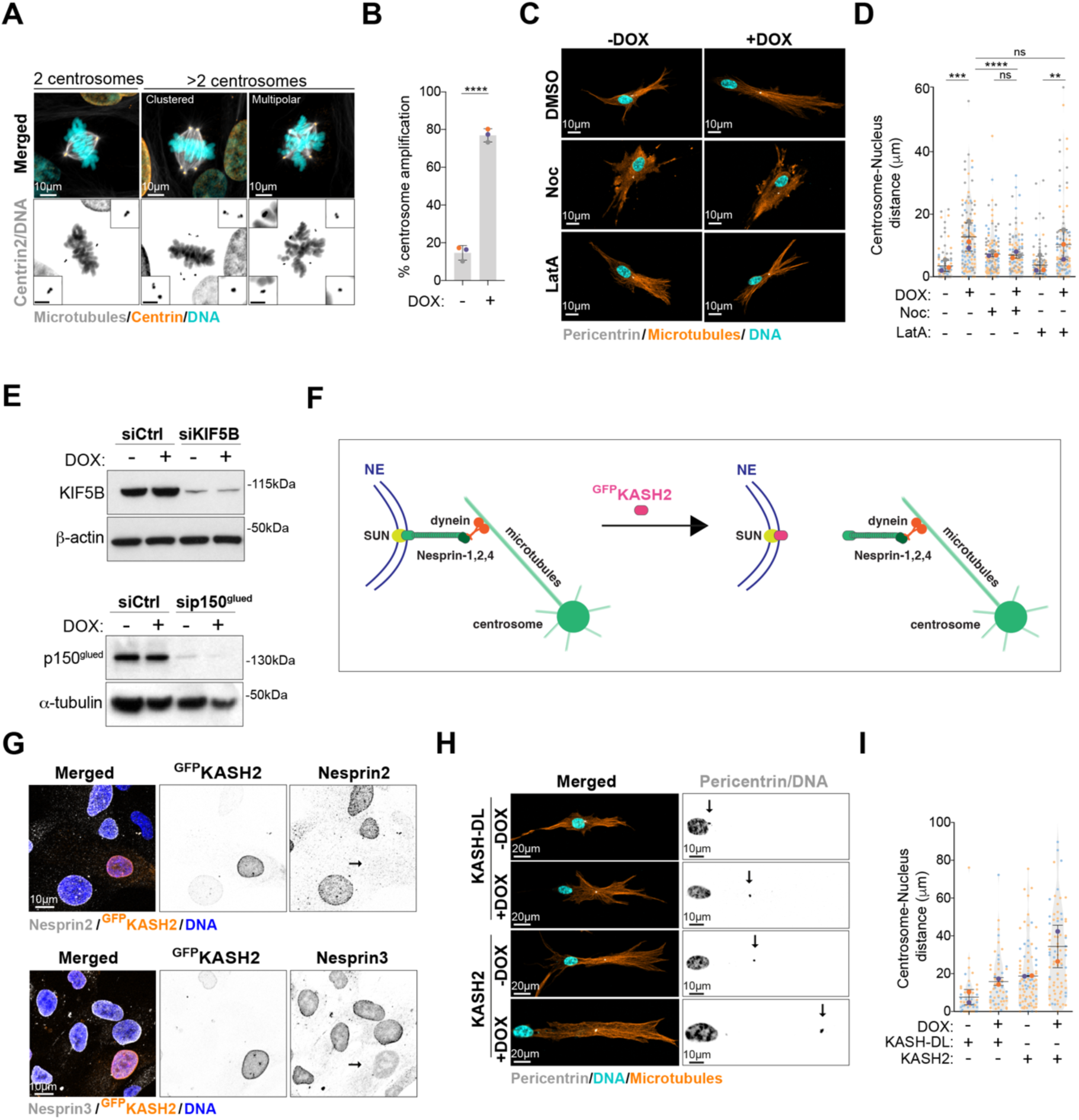
Microtubules mediate increased centrosome displacement in cells with extra centrosomes. **A.** Representative images of cells stained for centrosomes (Centrin2, cyan), microtubules (α-Tubulin, grey) and DNA (Hoechst, orange). Scale Bar: 10 µm; inset scale bar: 2 µm. **B.** Quantification of metaphase cells with extra centrosomes (n(-DOX)=337; n(+DOX)=339). **C.** Representative images of cells stained for centrosomes (Pericentrin, grey), microtubules (α-Tubulin, orange) and DNA (Hoechst, cyan) treated with Nocodazole (Noc, 10µM) or LatrunculinA (LatA, 100nM). Scale bar: 10 µm. **D.** Quantification of centrosome-nucleus distance (n(- DOX)=90; n(+DOX)=114; n(-DOX Noc)=112; n(+DOX Noc)=101; n(-DOX LatA)=110; + n(+DOX LatA)=108). **E.** Top panel; immunoblot of KIF5B and β-actin in cells after KIF5B siRNA for 48 h. Bottom panel; immunoblot of p150^glued^ and α-tubulin in cells after p150^glued^ siRNA for 48 h. **F.** Representative scheme of LINC complex and disruption following KASH2 overexpression. **G.** Top panel; representative images of cells stained for Nesprin2 (grey), ^GFP^KASH2 (orange) and DNA (Hoechst, blue). Scale bar: 10 µm. Bottom panel; representative images of cells stained for Nesprin3 (grey), ^GFP^KASH2 (orange) and DNA (Hoechst, blue). Scale bar: 10 µm. **H.** Representative images of cells stained for centrosomes (Pericentrin, grey), microtubules (α-Tubulin, orange) and DNA (Hoechst, cyan) overexpressing ^GFP^KASH2 or ^GFP^KASH-DL. Scale bar: 20 µm; inset scale bar: 10 µm. **I.** Quantification of centrosome-nucleus distance (*n*=2; number of cells ^GFP^KASH-DL(-DOX)=49; ^GFP^KASH-DL(+DOX)=55; ^GFP^KASH2(- DOX)=87; ^GFP^KASH2(+DOX)=82). For all graphics error bars represent mean +/- SD from three independent experiments. \*\**p* < *0.01, ***p* < *0.001, ****p* < *0.0001,* ns = not significant (*p* > *0.05*). The following statistic were applied: unpaired *t* test for graph in B and one-way ANOVA with Tukey’s post hoc test for graph in D.

**Figure EV2.**
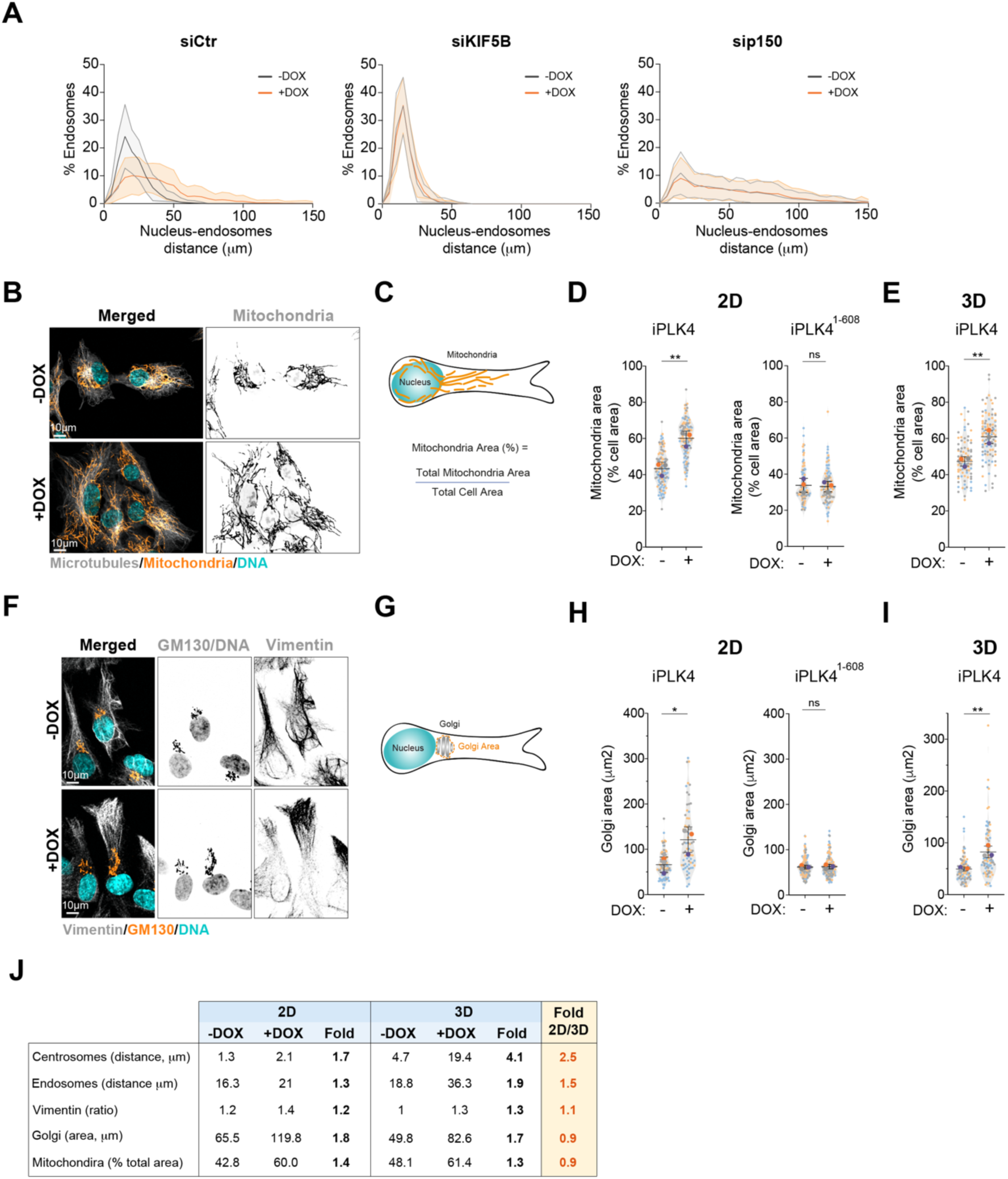
Increased mitochondria displacement and Golgi spreading in cells with extra centrosomes. **A.** Quantification of endosomes-nucleus distance upon depletion of KIF5B and p150 (n(-DOX siCtr)=84; n(+DOX siCtr)=81; n(-DOX siKIF5B)=84; n(+DOX siKIF5B)=83; n(-DOX sip150)=82; n(+DOX sip150)=83). **B.** Representative images of cells stained for mitochondria (MitoTracker, orange), microtubules (α-Tubulin, grey) and DNA (Hoechst, cyan). Scale bar: 10 µm. **C.** Representative scheme of mitochondria area quantification. **D.** Quantification of mitochondria area in cells plated in 2D upon PLK4 (Left panel; n(-DOX)=113; n(+DOX)=114) or PLK4^1-608^ overexpression (Right panel; n(- DOX)=95; n(+DOX)=98). **E.** Quantification of mitochondria area in cells plated in 3D (n(-DOX)=90; n(+DOX)=102). **F.** Representative images of cells stained for Golgi (GM130, orange), vimentin (grey) and DNA (Hoechst, cyan). Scale bar: 10 µm. **G.** Representative scheme of Golgi area quantification. **H.** Quantification of Golgi area upon induction of PLK4 (left panel; n(-DOX)=94; n(+DOX)=70) or PLK4^1-608^ overexpression (Right panel; n(-DOX)=146; n(+DOX)=133). **I.** Quantification of Golgi area in cells plated in 3D (n(-DOX)=91; n(+DOX)=83). **J.** Table summarizing the fold change between 2D and 3D conditions for the intracellular compartments analyzed. For all graphics error bars represent mean +/- SD from three independent experiments. \**p* < *0.05,* \*\**p* < *0.01,* ns = not significant (*p* > *0.05*). The following statistic were applied: unpaired *t* test for all graphs.

**Figure EV3.**
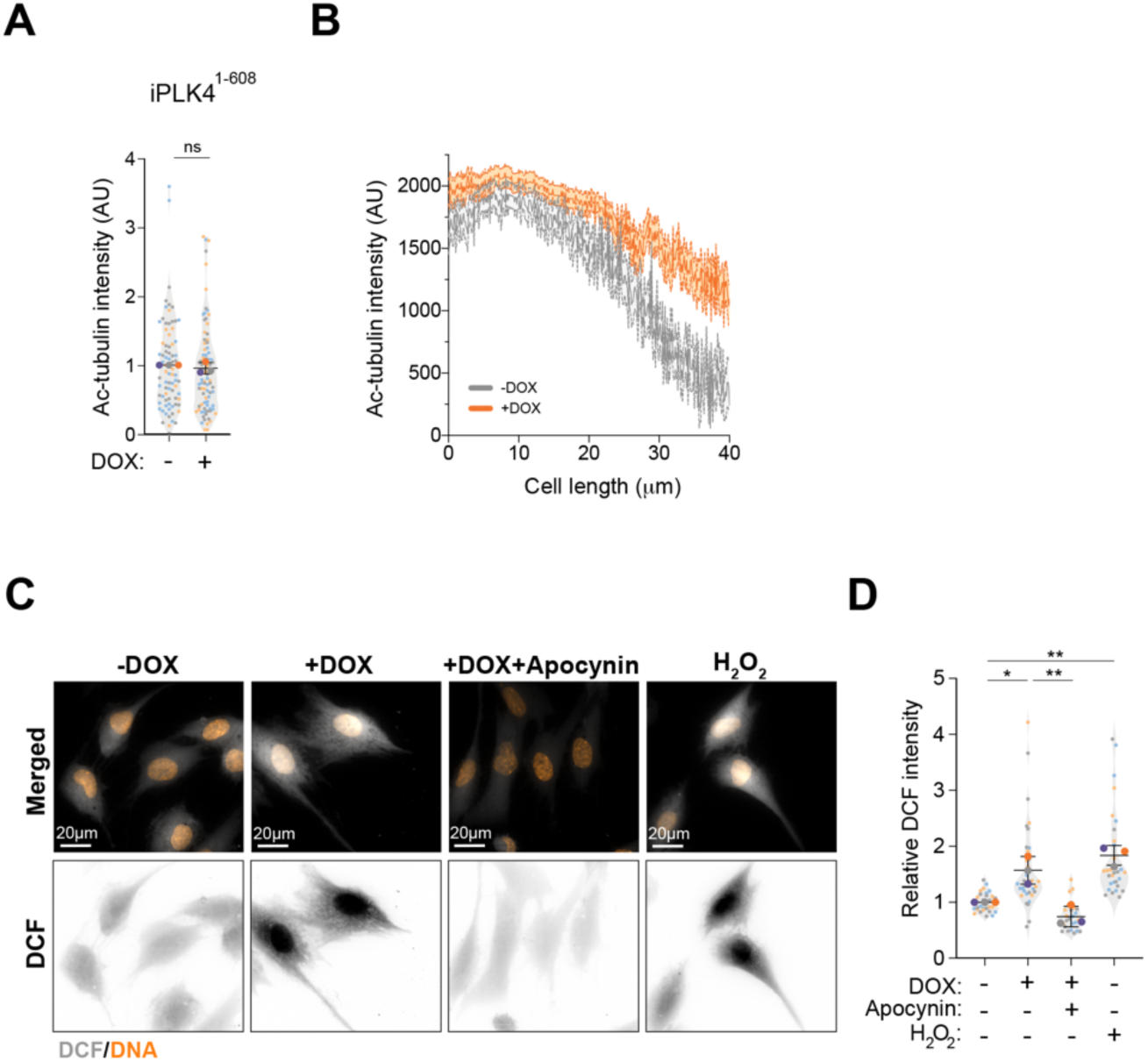
Centrosome amplification leads to stabilized microtubules and increased ROS levels. **A.** Quantification of acetylated tubulin fluorescence intensity in PLK4^1-608^ (n(-DOX)=78; n(+DOX)=85). **B.** Distribution of acetylated tubulin fluorescence intensity throughout the cell length. **C.** Representative images of cells stained for DNA (Hoechst, orange) and DCF (grey) treated with Apocynin (0.5mM) and H2O2 (75µM). Scale bar: 20 µm. **D.** Quantification of total DCF fluorescence intensity (n(-DOX)=30; n(+DOX)=36; n(+DOX Apocynin)=27; n(-DOX H2O2)=30). For all graphics error bars represent mean +/- SD from three independent experiments. \**p* < *0.05,* \*\**p* < *0.01,* ns = not significant (*p* > *0.05*). The following statistic were applied: unpaired *t* test for graphs in A and one-way ANOVA with Tukey’s post hoc test for graph in D.

**Figure EV4.**
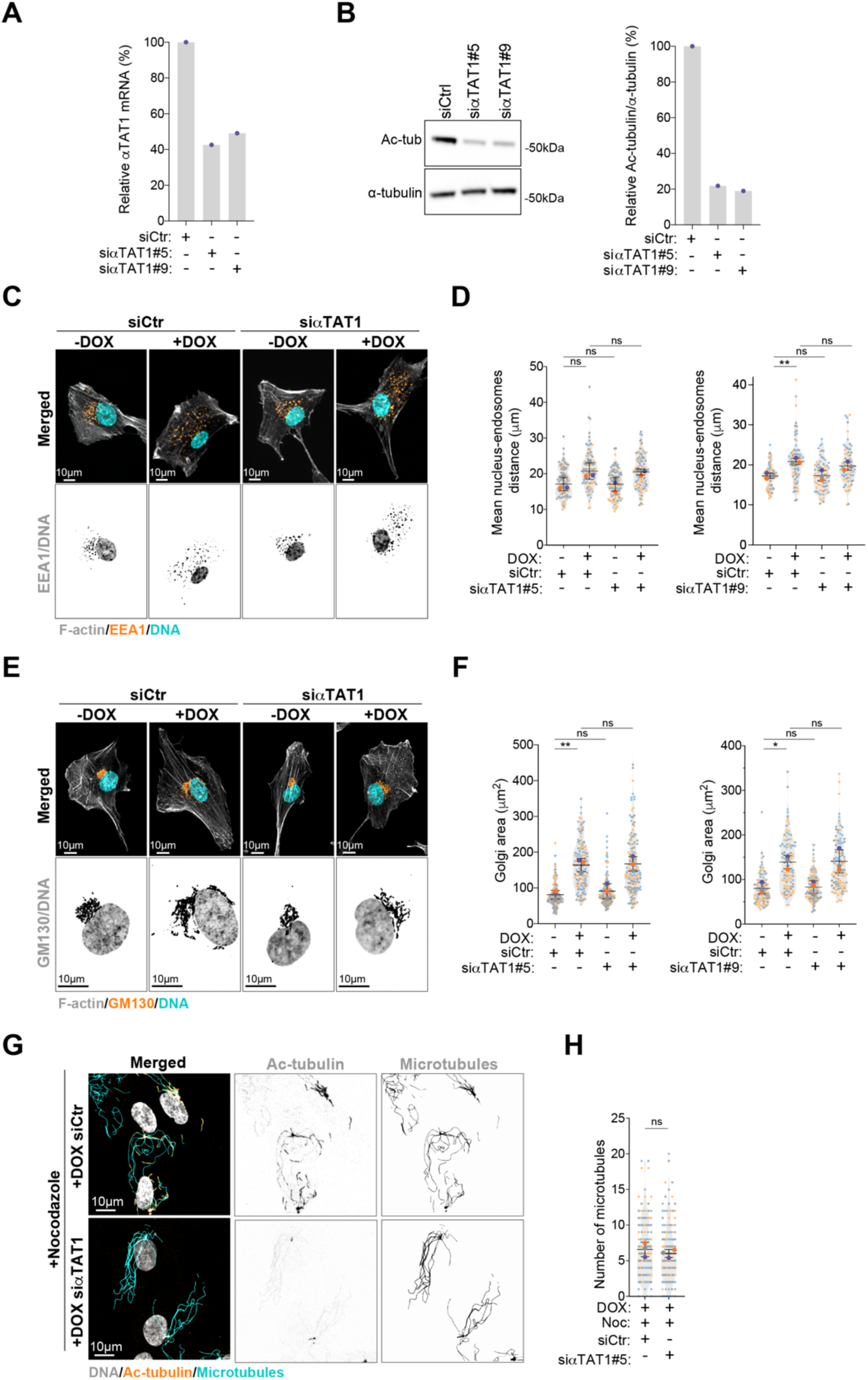
Endosomes displacement from the nucleus and increased Golgi area in cells with extra centrosomes do not rely on tubulin acetylation. **A.** Quantification of αTAT1 mRNA expression in cells treated with siRNA against αTAT1 (two independent siRNAs; #5 and #9). **B.** Left panel; immunoblot for α-tubulin and acetylated tubulin (Ac-tub) upon αTAT1 depletion (two independent siRNAs; #5 and #9). Right panel; percentage of acetylated tubulin levels per total α-tubulin. **C.** Representative images of cells stained for early endosomes (EEA1, orange), F-actin (phalloidin, grey) and DNA (Hoechst, cyan) upon αTAT1 depletion. Scale bar: 10 µm. **D.** Quantification of endosome-nucleus distance n(-DOX siCtr)=104; n(+DOX siCtr)=99; n(-DOX siαTAT1#5)=106; (+DOX siαTAT1#5)=101; n(-DOX siCtr)=76; n(+DOX siCtr)=79; n(-DOX siαTAT1#9)=77; (+DOX siαTAT1#9)=75). **E.** Representative images of cells stained for Golgi (GM130, orange), F-actin (phalloidin, grey) and DNA (Hoechst, cyan) upon αTAT1 depletion. Scale bar: 10 µm. **F.** Quantification of Golgi area (n(-DOX siCtr)=177; n(+DOX siCtr)=147; n(-DOX siαTAT1#5); n(+DOX siαTAT1#5)=156; n(-DOX siCtr)=103; n(+DOX siCtr)=103; n(-DOX siαTAT1#9)=128; n(+DOX siαTAT1#9)=109). **G.** Representative images of cells stained for microtubules (α-tubulin, cyan), tubulin acetylation (Ac-tubulin, orange) and DNA (Hoechst, grey) upon nocodazole treatment (Noc, 2µM). Scale bar: 10 µm. **H.** Quantification of microtubule numbers (n(+DOX siCtr +Noc)=145; n(+DOX siαTAT1 +Noc)=159). For all graphics error bars represent mean +/- SD from three independent experiments. \**p* < *0.05,* \*\**p* < *0.01,* ns = not significant (*p* > *0.05*). The following statistic were applied: one-way ANOVA with Tukey’s post hoc test for graphs in D and F and unpaired *t* test for graph in H.

**Figure EV5.**
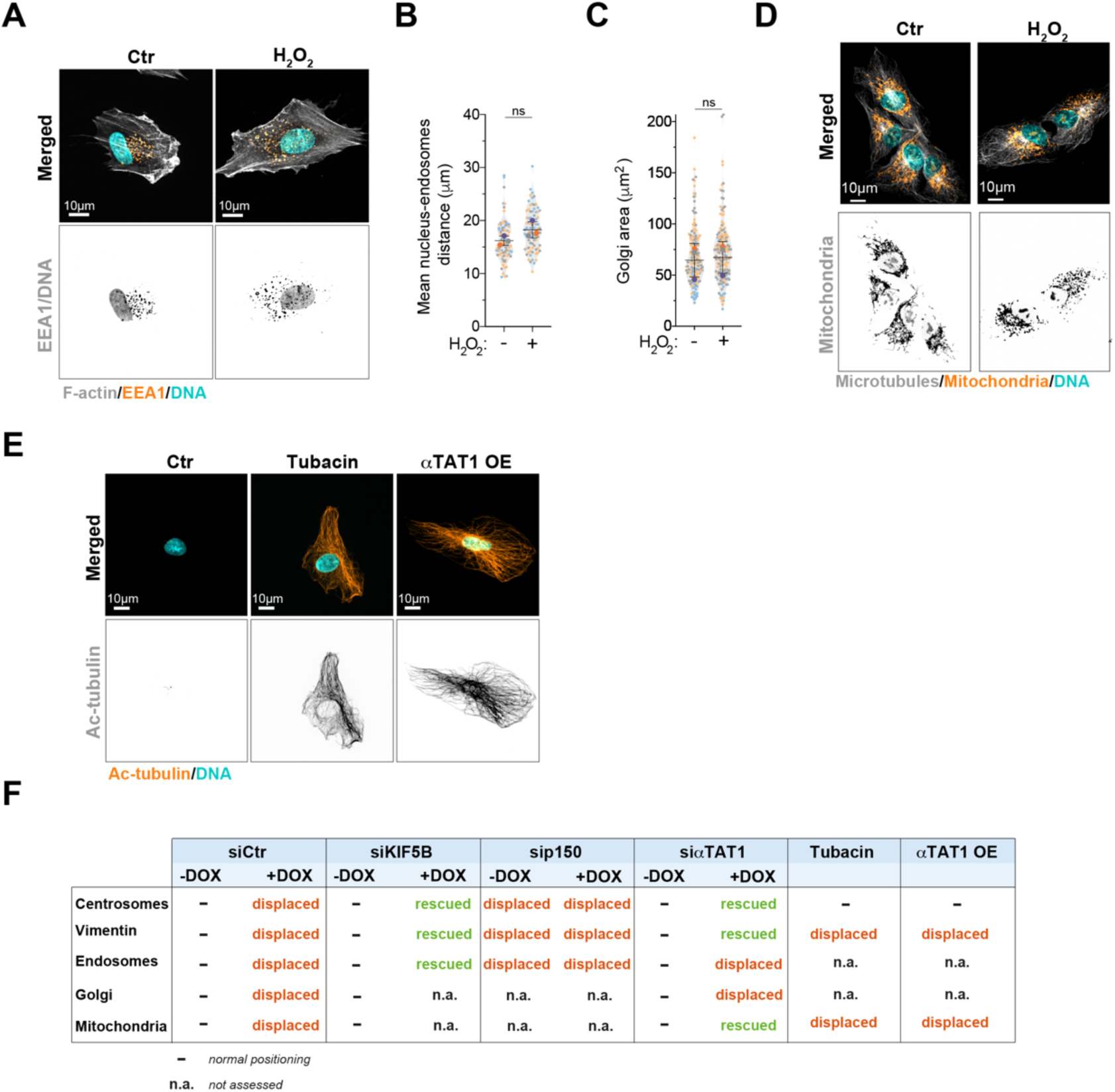
Endosomes and Golgi displacement in H2O2 treated cells. **A.** Representative images of cells stained for early endosomes (EEA1, orange), F-actin (phalloidin, grey) and DNA (Hoechst, cyan) treated with H2O2. Scale bar: 10 µm. **B.** Quantification of endosomes-nucleus distance (n(Ctr)=82; n(H2O2)=79). **C.** Quantification of Golgi area (n(Ctr)=151; n(H2O2)=157). **D.** Representative images of cells stained for mitochondria (MitoTracker, orange), microtubules (α-tubulin, grey) and DNA (Hoechst, cyan) treated with H2O2 (75µM). Scale bar: 10 µm. **E.** Representative images of cells stained for microtubules (α-tubulin, grey), tubulin acetylation (Ac-tubulin, orange) and DNA (Hoechst, cyan) in cells treated with Tubacin (5 µM) or overexpressing αTAT1-GFP (αTAT1 OE). Scale bar: 10 µm. **F.** Table summarizing the effect of different treatments on intracellular organization.

**Figure EV6.**
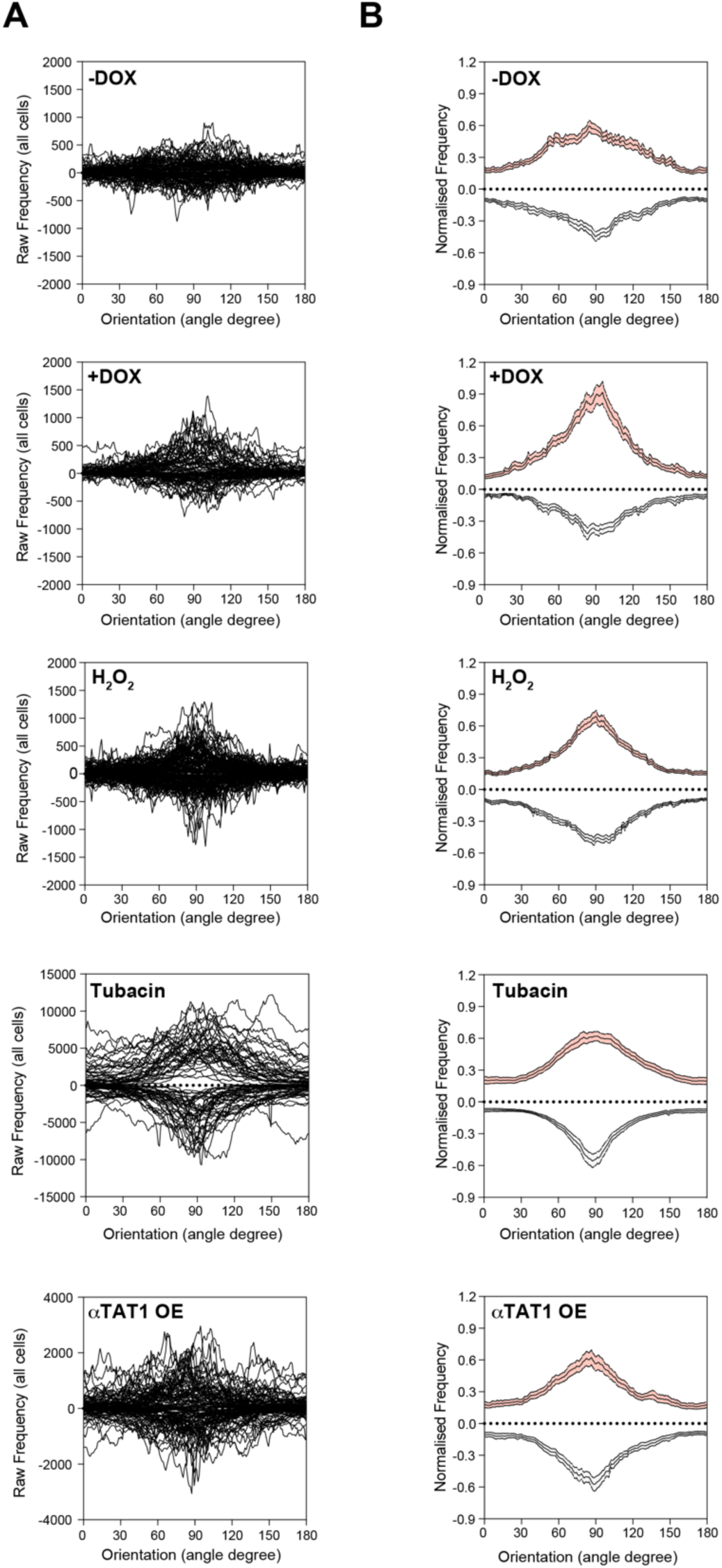
Distribution of acetylated microtubules in cells. **A.** Raw frequency of the distribution of acetylated microtubule orientation for each cell in cell front (+ values) and rear (- values). **B.** Normalized frequency of acetylated microtubule orientation (n(-DOX)=61; n(+DOX)=48; n(H2O2)=83; n(Tubacin)=39; n(αTAT1 OE)=63).

**Figure EV7.**
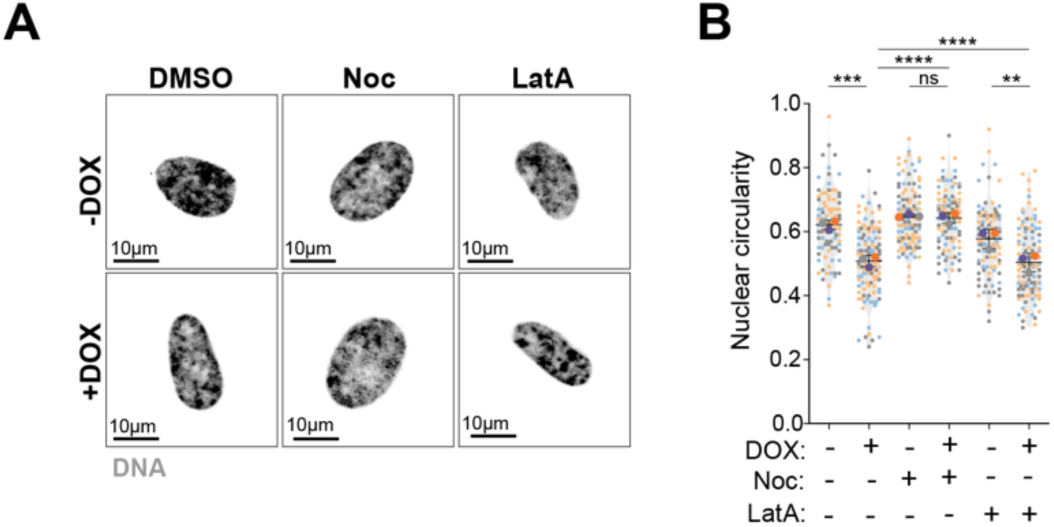
Microtubules control nucleus deformation in cells with extra centrosomes. **A.** Representative images of cell nucleus (DNA; Hoechst, grey) in cells treated with Nocodazole (Noc, 10µM) or LatrunculinA (LatA, 100nM). Scale bar: 10 µm. **B.** Quantification of nucleus aspect ratio (circularity) (n(-DOX)=90; n(+DOX)=114; n(-DOX Noc)=112; n(+DOX Noc)=101; n(-DOX LatA)=110; n(+DOX LatA)=108). For all graphics error bars represent mean +/- SD from three independent experiments. \*\**p* < *0.01, ***p* < *0.001, ****p* < *0.0001,* ns = not significant (*p* > *0.05*). The following statistic was applied: one-way ANOVA with Tukey’s post hoc test.

